# Dopamine neuron synaptic connectivity defines physiological striatal domains

**DOI:** 10.1101/2020.10.11.334961

**Authors:** Nao Chuhma, Soo Jung Oh, Stephen Rayport

**Affiliations:** Department of Molecular Therapeutics, New York State Psychiatric Institute, New York, NY 10032, USA; Department of Psychiatry, Columbia University, New York, NY 10032, USA

## Abstract

Dopamine neurons projecting to the striatum control movement, cognition, and motivation. They do so via slower, dopamine volume transmission and also via faster synaptic dopamine, glutamate and GABA transmission. To define the scope of these synaptic actions, we recorded dopamine neuron synaptic currents in the four major classes of striatal neurons. This revealed that dopaminergic and GABAergic synaptic actions are widespread; glutamatergic synaptic actions are robust in the medial nucleus accumbens and the anterolateral dorsal striatum, mediating fast and slow excitation, respectively. Dopamine neuron synaptic actions in cholinergic interneurons are the strongest and most complex, involving all three transmitters, their multiple receptors, and are the most regionally heterogeneous. The caudal striatum forms a single domain with overall weak dopamine neuron synaptic actions. This synaptic mapping reveals that dopamine neuron synaptic actions extend across the entire striatum, are regionally heterogeneous and organized in physiological domains, determined mainly by their excitatory actions.

## Introduction

Dopamine (DA) neurons connect principally with striatal (Str) neurons to control movement, cognition, and motivated behavior. Alterations in DA release underpin major neuropsychiatric disorders and are targeted for their pharmacotherapy. DA neurons in the ventral midbrain project to the Str topographically. More medial ventral tegmental area (VTA) neurons project to the ventral Str or nucleus accumbens (NAc), while laterally located substantia nigra (SN) neurons project to the dorsal Str or caudate-putamen (CPu) (Haber et al., 2000; Ikemoto, 2007). Genetically-specified intersectional strategies have revealed that distinct subclasses of DA neurons project differentially to Str subregions (Heymann et al., 2020; Kramer et al., 2018; Poulin et al., 2018; Wu et al., 2019). Despite regionally heterogeneous functions, the Str has a remarkably uniform cytoarchitecture, comprising about 95% spiny projection neurons (SPNs) and 5% interneurons. SPNs are GABAergic and equally divided in two types: SPNs expressing DA D1 receptors (D1R) that project directly to the ventral midbrain (direct-pathway SPNs; dSPNs), and SPNs expressing DA D2R that project to the ventral midbrain via the pallidum (indirect-pathway SPNs; iSPNs) (Gerfen and Surmeier, 2011). Str interneurons comprise cholinergic interneurons (ChIs) and GABAergic interneurons. While ChIs comprise a single type, GABAergic interneurons comprise multiple types; the most distinctive type are fast-spiking interneurons (FSIs) expressing parvalbumin (PV) (Tepper et al., 2018).

DA neurons signal via volume transmission (Sulzer et al., 2016), from varicosities with active zone-like release sites lacking postsynaptic elements (Liu and Kaeser, 2019). However, brain slice studies have also revealed that DA neurons have fast synaptic actions in some striatal regions mediated by three transmitters, DA, glutamate and GABA, and can directly excite or inhibit postsynaptic neurons. Glutamate cotransmission requires vesicular glutamate transporter VGLUT2 (Dal Bo et al., 2004; Hnasko et al., 2010), which is mainly expressed in DA neurons in the medial VTA (Morales and Margolis, 2017). These VGLUT2-expressing DA neurons project strongly to the medial shell of the NAc (Mingote et al., 2019; Poulin et al., 2018), where they invariably elicit excitatory synaptic responses via glutamate cotransmission (Stuber et al., 2010; Tecuapetla et al., 2010). DA neurons in the SN projecting to the CPu elicit synaptic DA (Chuhma et al., 2014), GABA (Tritsch et al., 2012) and glutamate (Chuhma et al., 2018) responses. GABA cotransmission requires vesicular monoamine transporter VMAT2 (Tritsch et al., 2012) and plasma membrane GABA transporter GAT1 (Tritsch et al., 2014). These sub-second synaptic responses mediated by the three neurotransmitters engage five receptors: ionotropic glutamate receptors (iGluR), metabotropic glutamate receptor 1 (mGluR1), DA D2R, DA D1/5 (D1-like) R, and GABA_A_R (Cai and Ford, 2018; Chuhma et al., 2014; Chuhma et al., 2018; Kim et al., 2015; Straub et al., 2014; Stuber et al., 2010; Sulzer et al., 1998; Tecuapetla et al., 2010; Tritsch et al., 2012; Wieland et al., 2014).

DA neuron synaptic transmission has only been examined in limited locations in the Str, so it has yet to be resolved whether DA neurons have widespread synaptic actions and how they are organized. Here we mapped DA neuron synaptic actions in identified Str neurons by recording direct synaptic responses from identified Str neurons. This has revealed that DA neuron synaptic actions extend across the entire Str and are organized in physiological domains.

## Results

### DA neuron synaptic currents evoked in principal striatal neuron types

To map DA neuron synaptic responses, we stimulated DA neuron axons impinging on the four principal Str neuron types, dSPNs, iSPNs, ChIs, and FSIs (**Fig. 1**). We used brain slices from triple mutant mice expressing channelrhodopsin 2 (ChR2) in DA neurons driven by the DA transporter (DAT^IREScre^; floxSTOP-ChR2 mice), and Str cell-type specific markers, D1-tdTomato for dSPNs, D2-enhanced green fluorescent protein (EGFP) for iSPNs, choline acetyltransferase (ChAT)-EGFP for ChIs, and PV-tdTomato for FSIs. While DA neurons in the dorsal raphe may also express ChR2, they do not project to the Str (Lin et al., 2020), so the evoked synaptic responses in the Str arise from ventral midbrain DA neurons. Since D2Rs are expressed in iSPNs, and also in ChIs (Lim et al., 2014), ChIs might express EGFP. However, in D2-EGFP mice only 16.5% of ChAT positive (ChAT^+^) neurons showed EGFP fluorescence (**Supplemental Fig. 1A, B**), which was significantly dimmer than in ChAT negative (ChAT^−^) neurons (Welch’s t-test, t = −10.3, p < 0.0001) (**Supplemental Fig. 1C**). ChAT^+^ neurons had a larger soma diameter than ChAT^−^ neurons regardless of EGFP fluorescence (one-way ANOVA, F_(2, 46)_ = 88.8, p < 0.0001) (**Supplemental Fig. 1D**). Therefore, in D2-EGFP mice smaller neurons with brighter EGFP fluorescence were identified as iSPNs.

**Fig 1.**
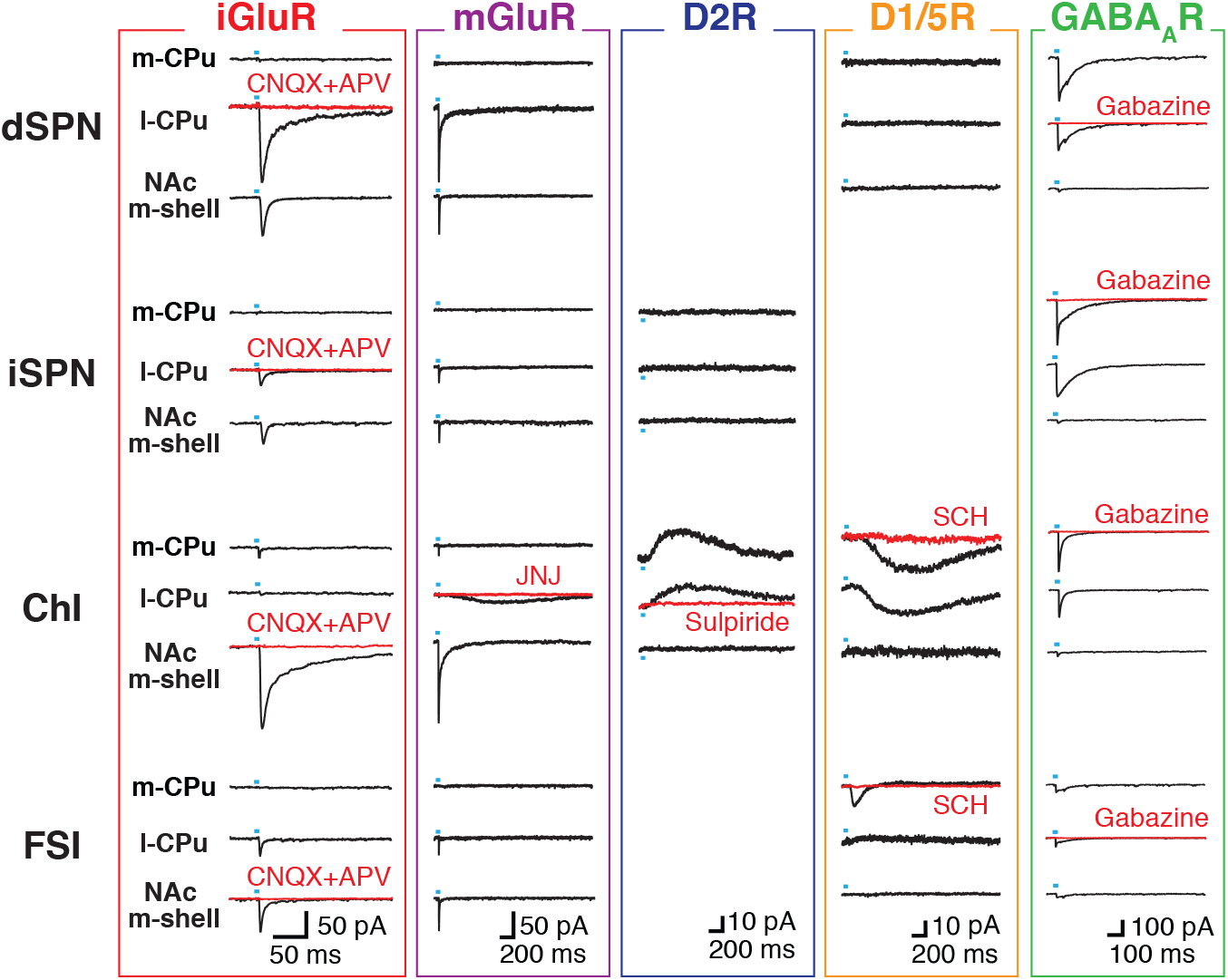
Dopamine-neuron evoked postsynaptic currents in striatal neurons. In DAT-iresCre;Ai32 mice with dopamine neurons made photoactivatable by expression of channelrhodopsin, pulse illumination (blue bar) of their terminals in striatal brain slices evoked synaptic currents in the four major types of striatal neurons, mediated by five different neurotransmitter receptors, that showed significant regional variation. Recorded striatal cell types were direct-pathway spiny projection neurons (dSPN), indirect-pathway SPN (iSPN), cholinergic interneurons (ChI), and fast-spiking GABAergic interneurons (FSI). Pharmacologically isolated postsynaptic currents (PSCs) were mediated by glutamate via iGluR and mGluR, dopamine via D2R and D1/5R, and GABA via GABA_A_R. The three recorded regions were the nucleus accumbens medial shell (NAc m-shell), the medial caudate-putamen (m-CPu) and lateral caudate-putamen (l-CPu). The same traces are shown for iGluR and mGluR, but on different time scales. PSCs were blocked by the respective antagonist for each transmitter receptor (red traces). D2R responses in dSPNs and FSIs, and D1/5R in iSPNs were not recorded, as the cells lack the corresponding receptors.

We photostimulated DA neuron axon terminals with single pulses of blue light, illuminating the entire field of view (5 msec duration, at 0.1 Hz), at maximum intensity, and recorded synaptic currents in Str neurons, clamping the membrane at −70 mV. Single-receptor type PSCs were pharmacologically isolated by bath application of a cocktail of antagonists to block other receptors: CNQX 40 µM and D-APV 100 µM for iGluRs, JNJ16259685 5 µM for mGluR1, SCH23390 10 µM for D1/5R, sulpiride 10 µM for D2R, and SR95531 (gabazine) 10 µM for GABA_A_R. In all recordings, nicotinic and muscarinic acetylcholine receptors (nAChRs, mAChRs) were blocked with mechamilamine (10 µM) and scopolamine (2 µM) to eliminate contributions from varying levels of ChI activity. A Cs^+^-based pipette solution was used for glutamate and GABA_A_ responses to improve space clamp, while a K^+^-based pipette solution was used for DA responses, so as not to block K^+^ channels. GABA_A_ responses were recorded with high Cl^−^ intracellular solution to obtain inward currents at the same holding potential as for the other responses.

The three DA neuron neurotransmitters elicited PSCs via five types of receptors: iGluRs, mGluR1, DA D1/5Rs and D2Rs, and GABA_A_Rs (**Fig. 1**). iGluR EPSCs and GABA_A_ IPSCs were observed in all four cell types, while mGluR EPSCs and D2 IPSCs were observed only in ChIs. D1/5 EPSCs were observed in ChIs and FSIs, but their time courses were different; FSI D1/5 EPSCs showed a faster time course (Rise time 10-90% peak: ChI 350.0 ± 14.7 ms, FSI 34.5 ± 2.1 ms, Welch’s t-test, t = 21.3, p < 0.0001; Width at 50% peak: ChI 807.0 ± 29.0 ms, FSI 94.9 ± 4.9 ms, t = 24.2, p < 0.0001; ChI n = 63 cells, FSI n = 26 cells). Thus, only ChIs showed all five types of PSCs. Mediation of each PSC was confirmed by application of the corresponding antagonist (**Fig. 2**; **Fig. 1**, red traces). All corresponding antagonist applications blocked the isolated PSCs (**Fig. 2**; statistics are reported in **Supplemental Table 1**). ChAT-GFP mice used for identification of ChIs overexpress vesicular ACh transporters (VAChT) (Crittenden et al., 2014; Nagy and Aubert, 2012), which might affect PSC measurements, with regional variation. Although AChRs were continuously blocked in all experiments, longer-term modulatory effects of VAChT overexpression might have a regionally different impact on size of PSCs. However, the size distribution of iGluR and mGluR EPSCs in NAc medial shell, medial CPu and lateral CPu was not significantly different in recordings made from ChAT-EGFP mice and non-ChAT-EGFP mice, in which ChIs were identified by size and physiology, confirmed by post-recording ChAT immunostaining (two-way ANOVA, genotype effects: iGluR F_(1, 57)_ = 0.14, p = 0.71; mGluR F_(1, 57)_ = 0.05, p = 0.82; interaction: iGluR F_(2, 57)_ = 0.16, p = 0.85; mGluR F_(2, 57)_ = 0.04, p = 0.96) (**Supplemental Fig. 2**). Dendrite lengths of ChIs and glutamate EPSC sizes were not correlated (linear regression, iGluR R^2^ = 0.0002, p = 0.96; mGluR R^2^ = 0.006, p = 0.76) (**Supplemental Fig. 3**), indicating that differences in the truncation of dendrites during slicing did not contribute to regional differences in PSCs.

**Fig 2.**
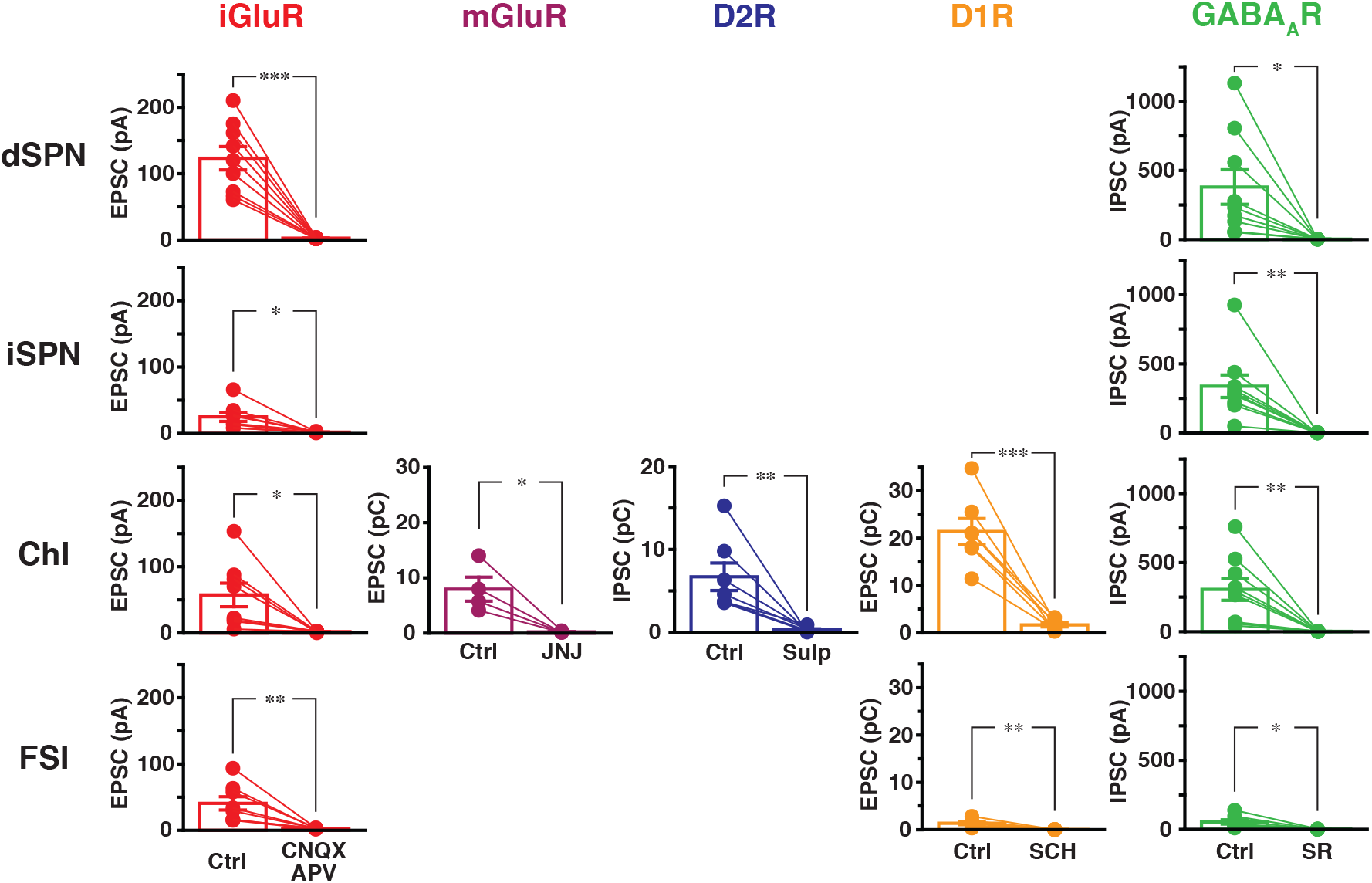
Receptor mediation of pharmacologically isolated PSCs. Pharmacologically isolated PSCs were blocked by the corresponding receptor antagonist. The antagonists and concentrations used were: iGluR (red) CNQX 40 µM and D-APV 100 µM, mGluR (purple) JNJ16259685 10 µM, D2R (blue) sulpiride 10 µM, D1/5R (orange) SCH23390 10 µM, GABA_A_R (green) SR95531 10 µM. PSC size measurements (dots) for control (Ctrl) and antagonist conditions for each cell are connected (lines). Bars show mean ± S.E.M. Results of paired t-tests are shown in Supplementary Table 1. *, **, and *** indicate p < 0.05, p < 0.01 and p < 0.001, respectively. See Table S1 for parameters of statistics.

We then recorded DA neuron synaptic responses in Str neurons throughout the Str, from a minimum of 100 cells for each response type. For iGluR and GABA_A_R PSCs, peak amplitudes were measured. For mGluR, D2R, D1/5R PSCs in ChIs charge transfer was measured in a 1.5 sec window. For the faster D1/5R PSCs in FSIs, we measured charge transfer in a 0.3 sec window. PSC sizes were color-scaled and plotted at the Str recording locations to reveal the spatial distribution (**Fig. 3**). iGluR EPSCs and GABA_A_ IPSCs were observed in all four cell types, but their distributions were different. iGluR EPSCs had two hotspots; the medial NAc in all four cell types and antero-lateral CPu in dSPNs, iSPNs, and FSIs. Averaged sizes of iGluR EPSCs, including cells without recognizable responses, were larger in dSPNs and ChIs, compared to iSPNs and FSIs (dSPN 24.6 ± 2.6 pA, n = 153 cells; iSPN 11.5 ± 1.2 pA, n = 149 cells; ChI 20.7 ± 2.9 pA, n = 133 cells; FSI 12.2 pA, n = 115 cells; one-way ANOVA, F_(3, 3546)_ = 8.3, p < 0.0001). GABA_A_ IPSCs did not show clear hotspots but were weaker in the more caudal Str and weak throughout the Str in FSIs (dSPN 158.1±17.0 pA, n = 138 cells; iSPN 119.0 ± 18.0 pA, n = 133 cells; ChI 211.8 ± 22.5 pA, n = 120 cells; FSI 31.8 ± 4.4 pA, n = 108 cells; one-way ANOVA, F_(3, 3495)_ = 16.9, p < 0.0001). mGluR EPSCs were observed only in the anterolateral CPu in ChIs. DA responses were not observed in SPNs, despite the high-expression of D1Rs in dSPN and D2Rs in iSPNs (Kreitzer, 2009; Surmeier et al., 2011). Indeed, charge transfers in SPNs measured as DA PSCs did not differ from baseline charge transfer recorded without photostimulation (two-sample Kolmogorov-Smirnov test; dSPN p = 0.48; iSPN p = 0.42), while D2R IPSCs in ChIs (positive control) showed a significant difference (p < 0.0001) (**Supplemental Fig. 4**). D2R IPSCs were observed widely in ChIs, stronger in the medial CPu and weaker in the medial NAc. D1/5R EPSCs were widely observed, both in ChIs and FSIs, without clear hotspots. D1/5R EPSC charge transfers in FSIs were smaller due to their faster time course.

**Fig 3.**
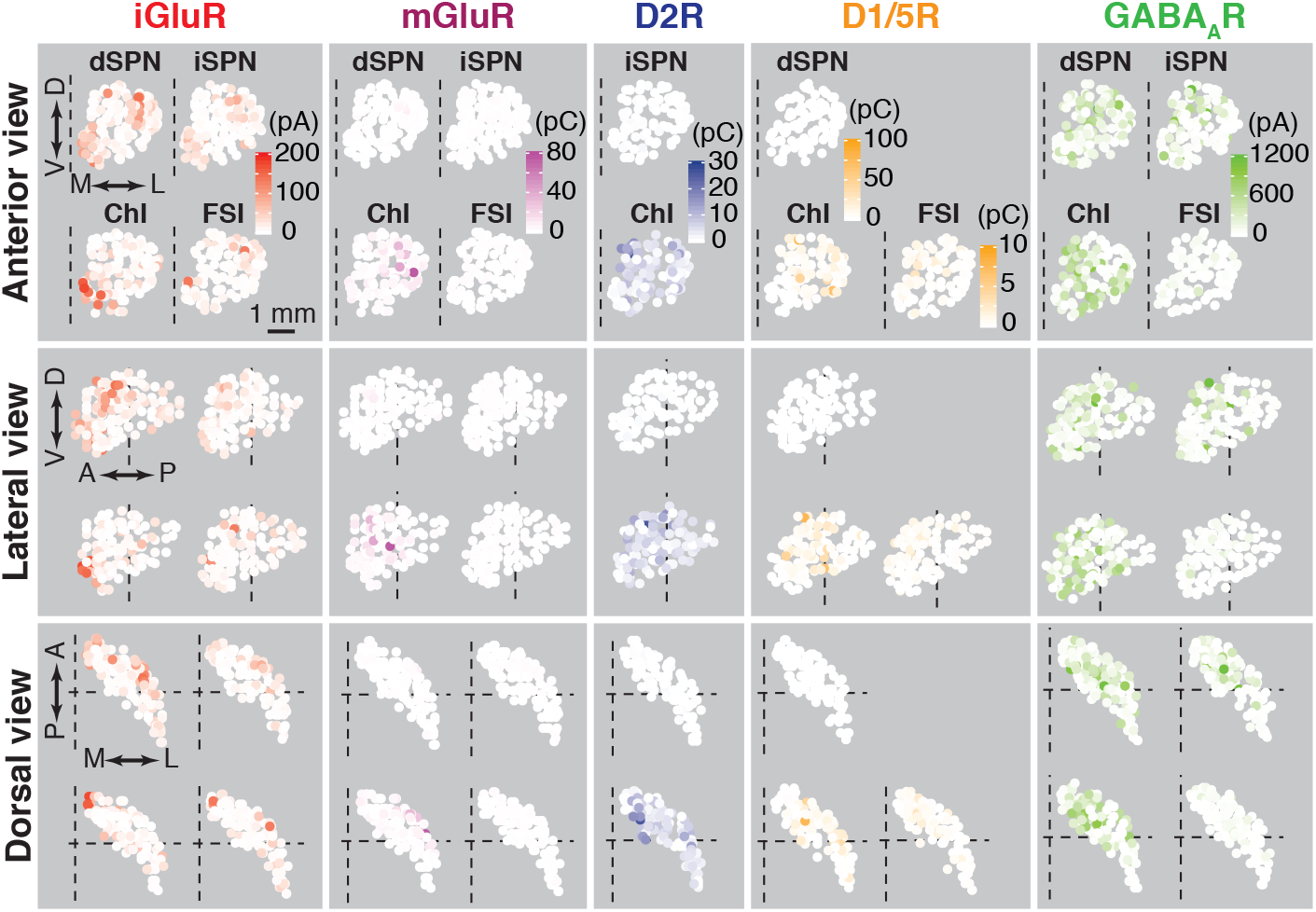
Distribution of dopamine-neuron evoked postsynaptic currents across the striatum. The size of PSCs, reported as amplitude for inotropic responses (iGluR, GABA_A_R, in pA) or charge transfer for metabotropic responses (mGluR, D2R, D1/5R, in pC), are plotted on a three-dimensional (3D) grid subtending the left striatum. Each recorded cell, indicated by a small sphere, is color coded by the size of the recorded PSC, with color intensity in red (iGluR), purple (mGluR), blue (D2R), orange (D1/5R), or green (GAB-AAR). Anterior (top), lateral (middle), and dorsal (bottom) views are shown; arrows indicate ventral (V), dorsal (D), medial (M), lateral (L), anterior (A), and posterior (P) directions. Dashed lines indicate bregma zero. D1/5R responses in FSIs are shown on a different color scale. PSCs were recorded in 1603 striatal cells: Glu dSPN 153, iSPN 149, ChI 133, FSI 115; D2 iSPN 101, ChI 121; D1/5 dSPN 104, ChI 121, FSI 107; GABAA dSPN 138, iSPN 133, ChI 120, FSI 108.

### Maps of DA neuron synaptic currents in the striatum

To generate continuous maps of PSC size, we interpolated individual measurements on a 150 µm cubic grid (4177 points in the unilateral Str) by kriging, a geostatistical method of estimation based on relatively limited sample numbers, using weighted averages (Matheron, 1963 1246-1266). This method assumes spatial autocorrelation; closer points are similar, while distant points are different (Bivand et al., 2013), and the weights for averaging are obtained based on sampled data. When semivariances of PSC sizes for any two locations were calculated and plotted against distance between them (experimental variogram), two points that were closer had smaller semivariance for all response types (**Supplemental Fig. 5**), showing spatial autocorrelation. Weights for averaging in kriging were obtained from the experimental variograms fitted by covariance functions (**Supplemental Fig. 5**; fitting parameters are in **Supplemental Table 2**).

We plotted estimated PSC sizes for each receptor type in the Str. Anterior, lateral, and dorsal views of surfaces of the 3D plots are shown in **Fig. 4**; coronal sections of the 3D plots are shown in **Fig. 5** for iGluR, mGluR, and D2R and in **Fig. 6** for D1/5R and GABA_A_R PSCs. The continuous maps showed response hotspots more clearly than the raw data plots. In iGluR PSC maps, all four cell types showed a hotspot in the medial NAc, although the intensity and distribution differed among cell types (**Fig.4**; **Fig. 5**). The dSPN, iSPN, and FSI hotspots were limited to the medial shell, while the ChI hotspot extended to almost the entire NAc. Another iGluR hotspot was seen in the anterolateral CPu for dSPNs, iSPNs, and FSIs, but not ChIs. Although FSIs showed both the medial NAc and antero-lateral CPu hotspots, the hotspots were smaller than those for the other cell types. Estimated iGluR responses in the caudal Str (caudal to bregma, bottom of **Fig. 5**) had recognizable sizes, but clear hotspots were not observed in all four cell types. In contrast to iGluR maps, mGluR responses were seen only in ChIs in one hotspot in the anterolateral CPu (**Fig. 4; Fig. 5**). DA D2R responses were widely observed throughout the Str, with a weak response spot in the NAc medial shell and a hotspot in the dorsomedial CPu (**Fig. 4; Fig. 5**). Despite the lack of clear hotspots for D1/5R or GABA_A_R responses in the raw data plots (**Fig. 3**), the estimated response maps showed some spatial patterns (**Fig. 4**; **Fig. 6**). D1/5R responses in ChIs formed an oblong hotspot in the CPu near bregma, and a second hotspot in the middle of the anterior CPu (**Fig. 4**). Coronal sections revealed that the oblong hotspot and the anterior CPu hotspots were connected, and also revealed weak response areas in the medial CPu and NAc (**Fig. 6**). In FSIs, D1/5R responses were more prominent in the medial CPu (**Fig. 6**). Overall GABA_A_R responses were weaker in FSIs compared to the other three cell types. In SPNs and ChIs, GABA_A_R responses tended to be larger in the anterior Str, and weaker in the most caudal Str (< −1.0 mm from bregma) (**Fig. 6**). In SPNs, the middle part of the anterior Str showed larger GABA_A_ responses, and the anterolateral CPu showed weaker responses. In ChIs, the medial NAc and anterior dorsolateral CPu showed weaker GABA_A_ responses, as well as the caudal-most Str.

**Fig 4.**
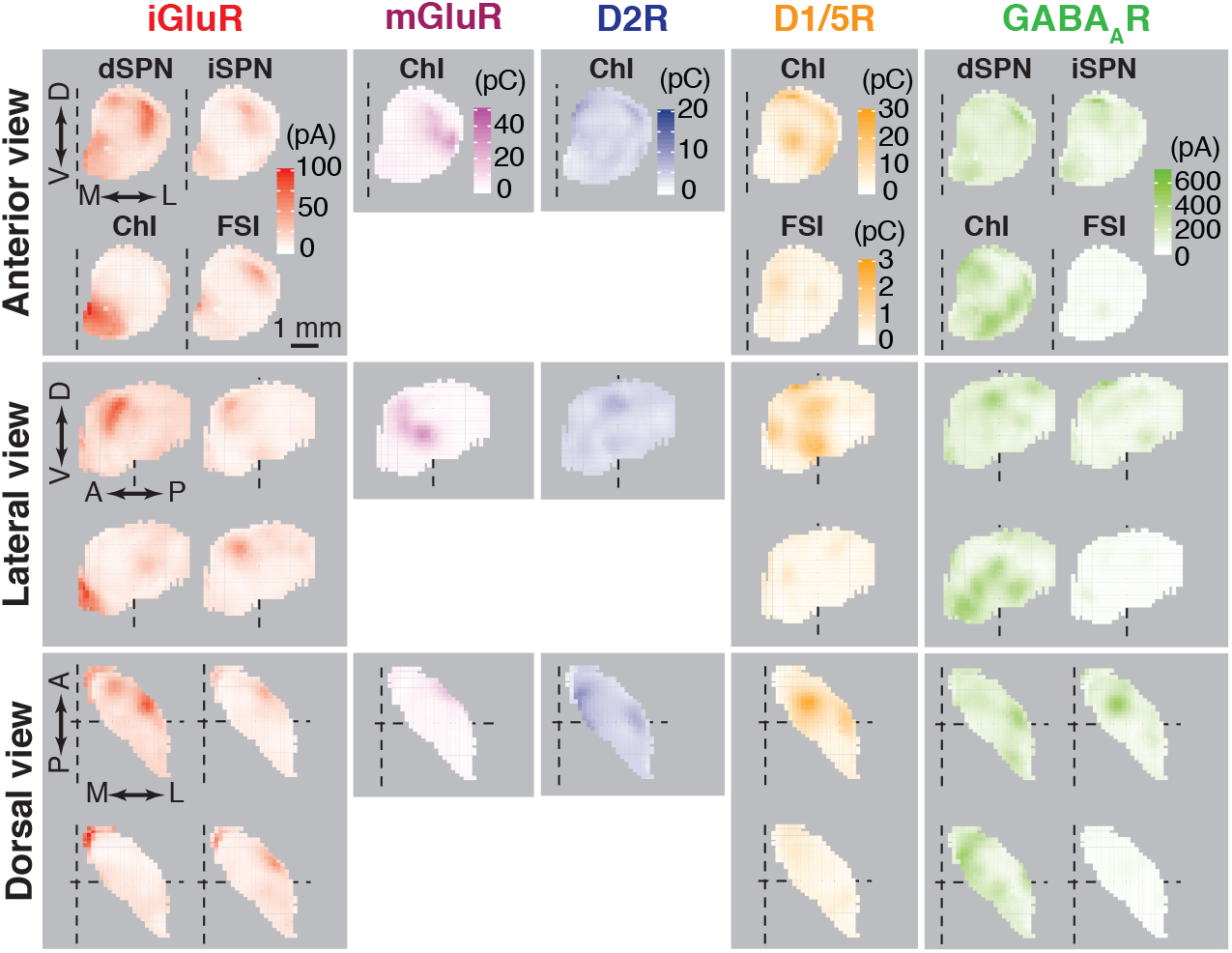
Maps of estimated dopamine-neuron evoked postsynaptic currents in 3D. Maps of estimated PSC size on a ∼150 µm 3D grid were made based on recorded PSC data using geostatistical methods. Only cell type/receptor recordings with significant responses were mapped. Maps maintain the same color scheme and layout as Fig. 3.

**Fig 5.**
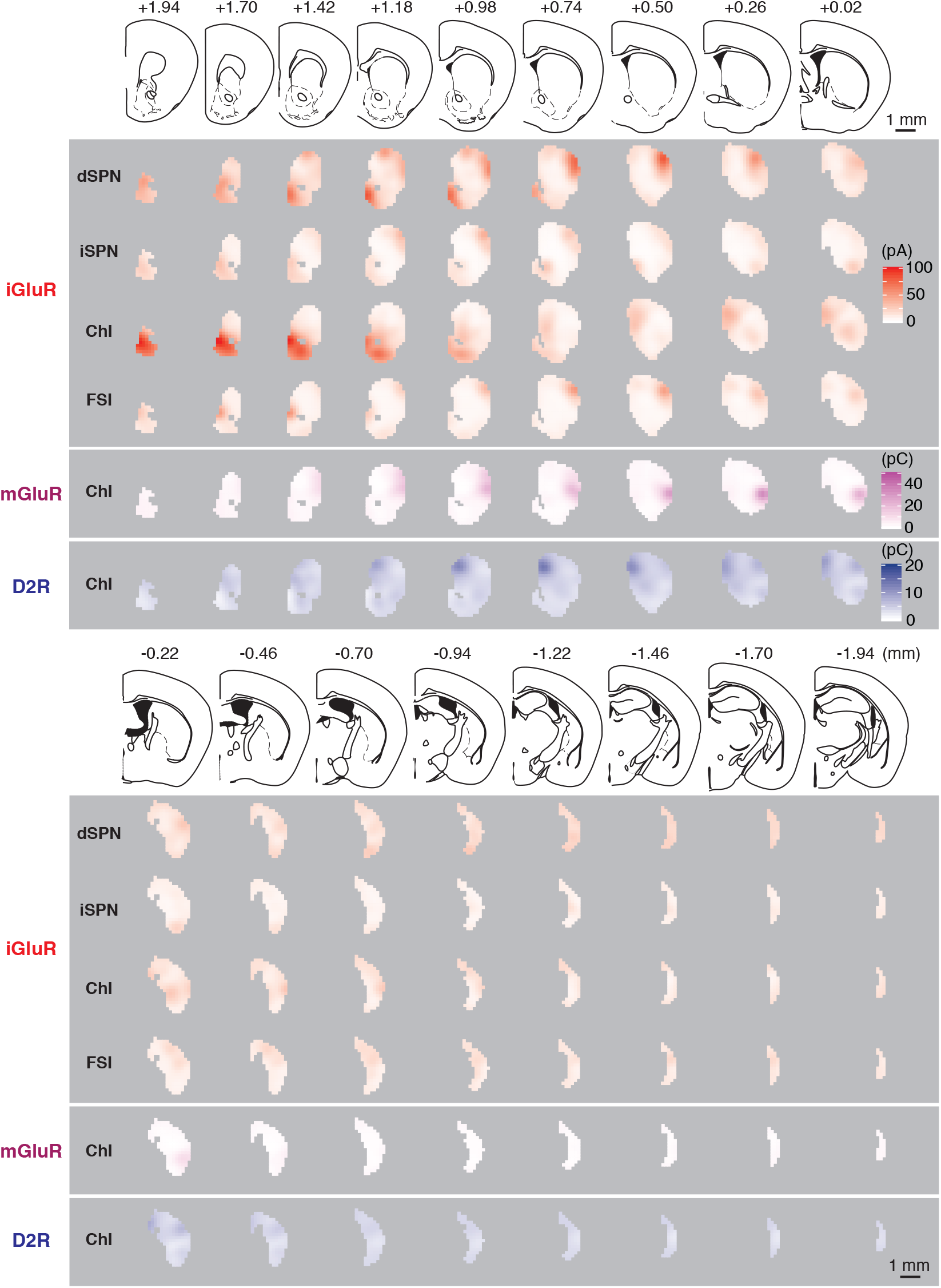
Maps of estimated dopamine-neuron evoked iGluR, mGluR, and D2R PSCs in coronal sections. Estimated iGluR (red), mGluR (purple), and D2R (blue) PSC sizes in the Str in coronal sections every ∼ 250 µm are shown, corresponding to every other left-sided slice in mouse atlas. Anterior-posterior location from bregma and unilateral brain atlas sections are shown above.

**Fig 6.**
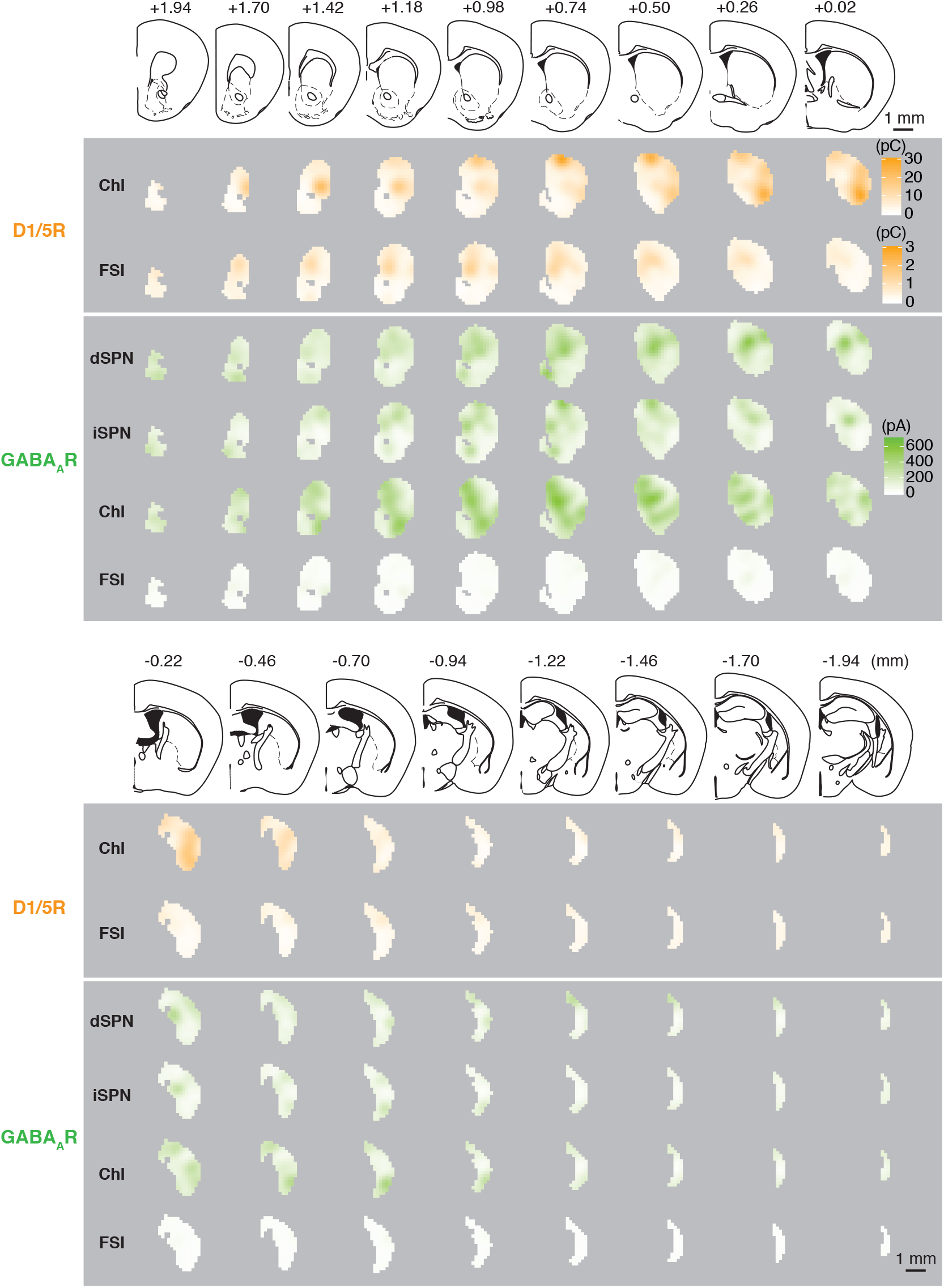
Maps of estimated dopamine-neuron evoked D1/5R and GABAAR PSCs in coronal sections. Estimated D1/5R (orange) and GABA_A_R (green) PSC sizes in the Str in coronal sections are shown with the same scheme as Fig 5.

### DA neuron synaptic connections define physiological striatal domains

To identify Str subregions based on synaptic connectivity for all observed transmitter responses in each Str cell type, we performed a cluster analysis for each cell type (**Fig. 7**) and for each transmitter (**Fig. 8**), using hierarchical clustering with spatial constraint (Chavent et al., 2018), mixing location and data (response size) matrices. PSC sizes were normalized to the maximum and minimum values of each map before clustering. Optimal numbers of clusters were determined by the silhouette method (Rousseeuw, 1987). In each cluster, we calculated mean and standard deviation (SD) of each response type, and the number of grid points as an estimate of cluster size (**Supplemental Table 3; Supplemental Table 4**). We also calculated cluster-separation F values, the ratio of between-cluster variance to within-cluster variance (Calinski and Harabasz, 1974), for each type of response to identify principal determinants of the clusters (**Supplemental Table 3; Supplemental Table 4**). Larger F values indicate better separation of clusters, and that the response type contributes more to the clustering.

**Fig 7.**
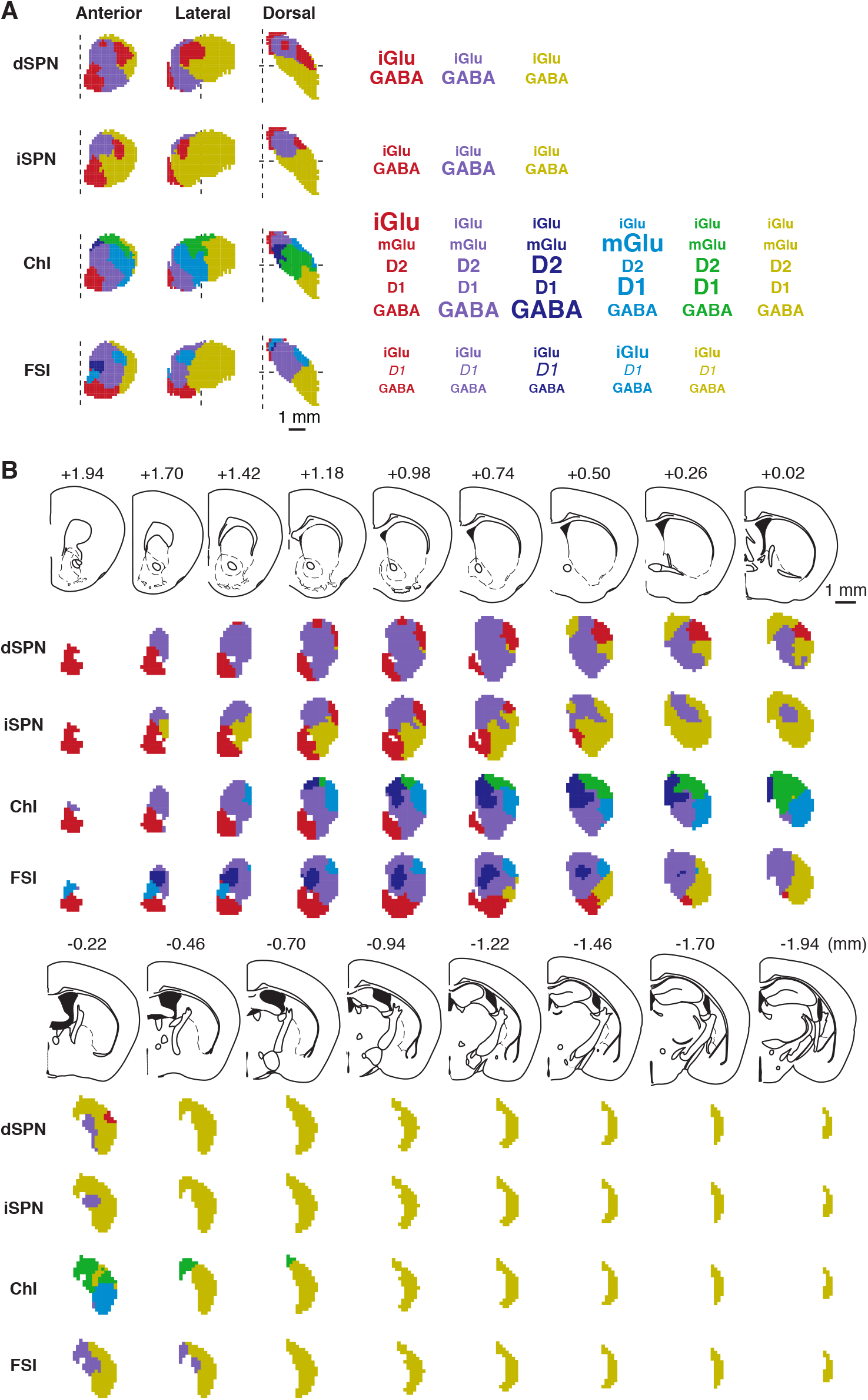
Domains of dopamine-neuron evoked PSCs by Str cell-type. Clusters by striatal cell type based on estimated PSC sizes are shown in anterior, lateral, and dorsal 3D views (**A**) and in coronal sections (**B**). Right panel in **A** shows mean PSC size for each cluster scaled by font size relative to the maximum response for each PSC type. D1/5R PSCs in FSIs are italicized to indicate that they were measured on a different scale than in ChIs.

**Fig 8.**
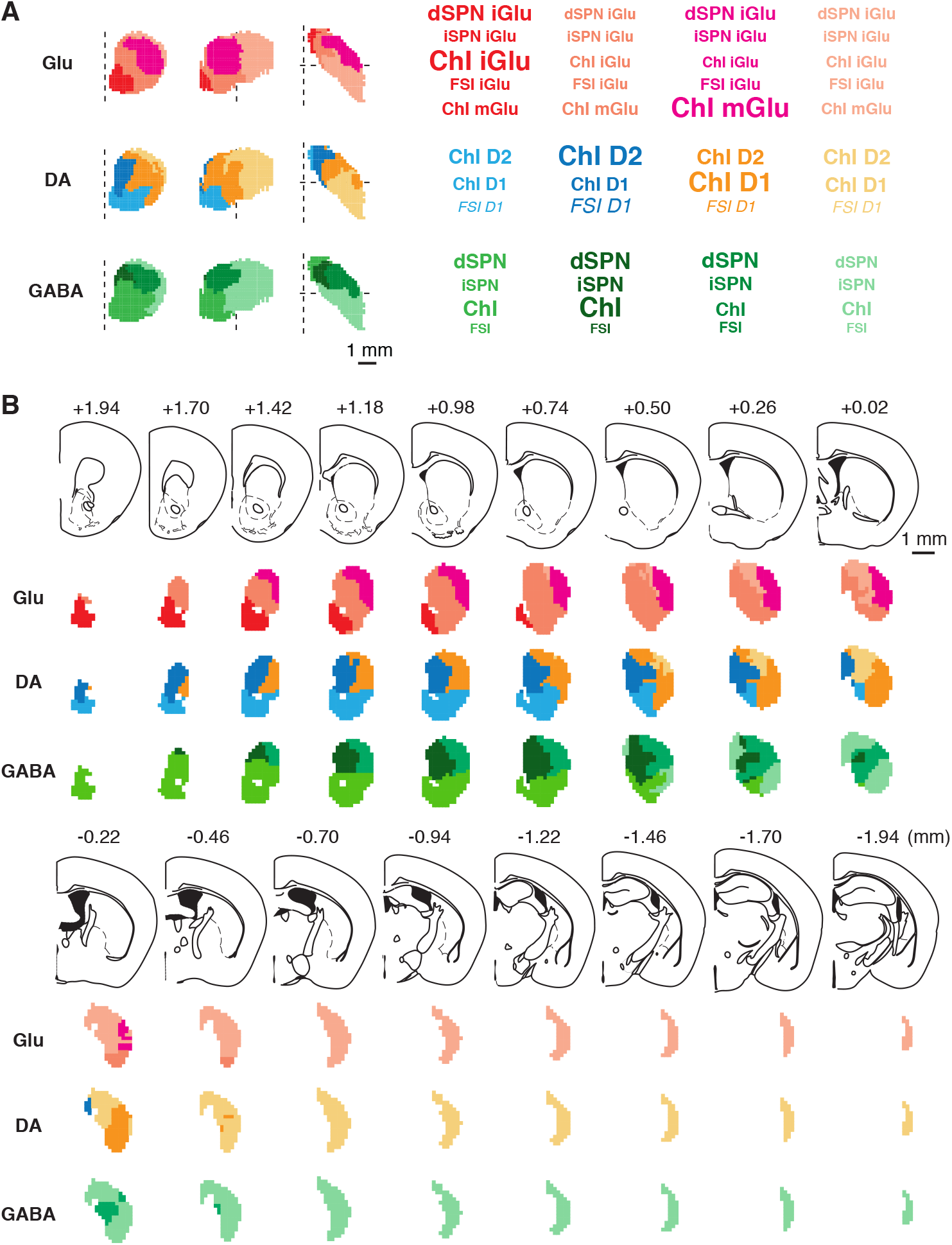
Domains of dopamine-neuron evoked PSCs by neurotransmitter. Clusters by DA neuron neurotransmitter are shown in anterior, lateral, and dorsal views (**A**) and in coronal sections (**B**). Right panel shows mean PSC size for each cluster scaled by font size relative to the maximum response for each PSC type, with the same scheme as Fig 7.

In cell-type clusters for both types of SPNs, the optimal number of clusters was three: the medial NAc and anterolateral CPu formed one cluster with the strongest iGluR PSCs, a second cluster in the posterior CPu where both iGluR and GABA_A_R PSCs were weak, and a third cluster that comprised the rest of the Str (**Fig. 7**). However, the posterior CPu cluster in iSPNs extended more anteriorly than in dSPNs, over most of the lateral CPu. In dSPNs, the iGluR F was substantially larger than the GABA_A_R F, suggesting that clusters were determined more by iGluR responses (**Supplemental Table 3**). In iSPNs, the GABA_A_R F was larger, but the difference between iGluR and GABA_A_R was not as large as in dSPNs, suggesting that both iGluR and GABA_A_R responses contributed equally to the clustering (**Supplemental Table 3**). In ChIs, all five types of responses were observed and the optimal number of clusters was six (**Fig. 7**). The medial NAc cluster and anterolateral CPu cluster were characterized by stronger excitatory responses; iGluR in the former, and mGluR and D1/5R in the latter. The anteromedial CPu formed a distinct cluster, where inhibition was dominant due to the strongest D2R and GABA_A_R responses. The caudal CPu showed generally small responses, as observed in SPNs. iGluR and GABA_A_R responses in ChIs were the largest among those in the four cell types (**Supplemental Table 3**). The most prominent determinant of clusters in ChIs was iGluR followed by D2R and GABA_A_R responses (**Supplemental Table 3**). In FSIs, although all responses were small, there were five clusters: a dorsomedial NAc and anterolateral CPu cluster with larger iGluR and GABA_A_R PSCs, an anterior dorsomedial CPu cluster with the strongest D1/5R responses, and a caudal CPu cluster with weak responses (**Fig. 7**). The strongest determinant of FSI clusters was D1/5R responses (**Supplemental Table 3**).

Transmitter clusters confirmed glutamate hotspots observed in cell-type clusters (**Fig. 8**). When clustering was done with all types of glutamate responses in all cell types (glutamate clusters), the optimal number of clusters was four (**Fig. 8**). Among them, two clusters were highlighted; the medial NAc characterized by iGluR responses in ChIs and dSPNs, and the anterolateral CPu characterized by mGluR responses in ChIs and iGluR responses in dSPNs. The clusters were determined by ChI iGluR, dSPN iGluR, and ChI mGluR PSCs, according to F values (**Supplemental Table 4**). In clustering of DA responses, the optimal number of clusters was four: medial NAc and posterior CPu clusters with weaker DA responses, a medial CPu cluster with prominent D2R responses, and an anterolateral CPu cluster with prominent D1/5R responses (**Fig. 8**). DA clusters were determined mostly by D1/5R responses, suggesting that D2R responses are distributed more widely and uniformly and contribute less to the clustering (**Supplemental Table 4**). The optimal number of GABA clusters was four, and GABA clusters showed the same tendency in all cell types: strongest in the antero-dorsomedial CPu, and weakest in the posterior CPu (**Fig. 8**). Clusters were determined almost equally by dSPN, iSPN, and ChI, but not by FSIs (**Supplemental Table 4**). Transmitter clusters supported another observation in the cell-type clustering; in addition to the two glutamate hotspots, the posterior CPu formed a single cluster with universally weak DA neuron synaptic responses. Taken together, clusters were predominantly determined by glutamate and D1/5R responses, compared to D2R or GABA_A_R responses. Both glutamate and DA D1/5 receptors mediate excitatory responses, indicatinging that DA neuron physiological domains in the Str are determined mainly by their excitatory synaptic actions.

## Discussion

Over time, DA neurons have gone from a homogeneous group of neurons recognized by their monoamine content and tyrosine hydroxylase expression, to subpopulations receiving differential inputs, projecting differentially, with diversity in their activity patterns, gene expression, cotransmission and behavioral functions. This heterogeneity appears to extend to DA neuron synaptic actions in the Str. However, the direct synaptic actions of DA neurons have only been identified in some DA neuron projections, in limited Str regions. Here we have mapped DA neuron synaptic actions in the Str, revealing that DA neurons have fast synaptic actions extending across the Str. SPNs show only iGluR and GABA_A_R responses, despite high expression of DA receptors. FSIs show D1/5R responses in addition to iGluR and GABA_A_R responses. Among the four principal types of Str neurons, only ChIs show responses to the three DA neuron neurotransmitters, mediated by five types of postsynaptic receptors. Maps of PSC size identify the most robust connections in the medial NAc for iGluR responses, anterolateral CPu for iGluR and mGluR responses, and medial CPu for D2R responses. Cell type clustering and transmitter clustering reveal discrete physiological domains: the medial NAc and anterolateral CPu as excitatory-dominant clusters, and the caudal CPu as a single cluster with universally weak synaptic connections. Inhibitory responses are distributed more broadly in the Str, so clusters are determined mainly by excitatory glutamate and D1/5R actions.

### DA neuron synaptic transmission in the Str

Ventral midbrain DA neurons have two temporal modes of transmission; slower volume transmission (Sulzer et al., 2016), and faster, point-to-point synaptic transmission. DA neuron DA synaptic transmission was first revealed in recordings of dendrodendritic connections between DA neurons in the ventral midbrain (Beckstead et al., 2004). In the Str, DA neuron glutamate connections were revealed with electrical stimulation of DA neuron axons (Chuhma et al., 2004) and optogenetic stimulation of DA neuron terminals impinging on SPNs in the NAc (Stuber et al., 2010; Tecuapetla et al., 2010). DA neurons made stronger iGluR-mediated connections to ChIs in the NAc m-shell, and direct DA D2R-mediated connections in the dStr (Chuhma et al., 2014). The present results show that fast DA neuron synaptic signaling spans the entire Str. DA neuron synaptic actions are not only faster, but also show cell-type specificity and convey discrete information, with regional heterogeneity due to blending of different types of responses.

In addition to fast PSCs mediated by ion channel-type receptors, i.e. iGluR and GABA_A_R, G-protein coupled receptors also mediate faster signals in Str interneurons. Stimulation of DA neuron axons elicits D2R IPSCs, D1/5 EPSCs, and mGluR EPSCs in ChIs (Cai and Ford, 2018; Chuhma et al., 2014; Chuhma et al., 2018; Straub et al., 2014; Wieland et al., 2014). While Str neurons express GABA_B_ receptors, GABA_B_-mediated PSCs were not observed in any Str cell types; GABA_B_ receptors are localized presynaptically (Lacey et al., 2005) and inhibit SPN collaterals (Shi and Rayport, 1994), and. D1/5R mediated depolarization and inward currents were observed in FSIs with bath application of D1 agonist (Centonze et al., 2003). However, perfusion is not the same as synaptic transmission, so it was not known whether action potential-dependent DA release would elicit D1/5R EPSCs in FSIs. We found that DA neurons elicit D1/5 EPSCs in FSIs, while SPNs show only iGluR and GABA_A_R PSCs despite their high DA receptor expression (Kreitzer, 2009; Surmeier et al., 2011). iSPNs show D2R IPSCs following transfection with G-protein coupled inward rectifier K^+^ (GIRK2) channels (Marcott et al., 2014), so the observation of fast DA PSCs in interneurons appears to depend on the expression of effector channels mediating currents, rather than DA receptors alone. Indeed, the D2R response in GIRK2 transfected iSPNs is slower in its latency and time course than that in ChIs, suggesting that other factors determine the time course of DA synaptic transmission such as the distance from DA release sites to DA receptors.

We identified DA neuron synaptic domains by cell-type cluster and transmitter-type cluster analysis. Both types of clusters were mainly determined by excitatory synaptic actions, i.e. glutamate and D1/5R responses, and mostly located in the anterior CPu (anterior to bregma) and NAc. The posterior CPu (posterior to bregma) formed its own cluster for all four cell types, with comparatively weak connections that showed little variation, indicating that most DA neuron signaling in the posterior CPu is via volume transmission. Although hotspots of glutamate cotransmission identified in this study correspond to locations of dense projections of glutamate cotransmitting DA neurons identified by intersectional expression of marker proteins (Chuhma et al., 2018; Mingote et al., 2019; Poulin et al., 2018), dense projections are also observed in the most caudal Str (Poulin et al., 2018), which belongs to the posterior CPu cluster, with weak synaptic connections. This discrepancy could be due to fewer release sites per projection fiber in the most caudal Str, as denser projections do not necessarily release more transmitters. Another possibility is that the projections in the caudal CPu synapse on other cell types, i.e. other types of GABA interneurons or glia. This is a potential limitation of the present mapping study, but it does emphasize further that anatomical evidence of connectivity does not translate directly to physiological measures of connectivity.

### Functional implications

DA neurons target ChIs preferentially and elicit the strongest responses, in contrast to the universally weak responses in FSIs. Considering their shallower resting membrane potential and spontaneous firing (Kawaguchi, 1993; Kreitzer, 2009), ChIs in the medial NAc and anterolateral CPu are the most easily excited by DA neuron synaptic inputs. Indeed, excitatory inputs from DA neurons can drive ChIs to fire, despite substantial GABA_A_ input (Chuhma et al., 2014; Chuhma et al., 2018; Straub et al., 2014). Although the two subregions are excited by DA neuron firing, the timing of the excitation differs due to differences in receptor mediation, iGluR versus mGluR and D1/5R. In other regions, DA neuron synaptic inputs appear to have inhibitory effects, pausing ChIs in the medial CPu via D2R and GABA_A_ receptors. Thus, DA neurons appear to use ChIs as hubs to control the Str circuitry with profoundly different synaptic effects in different Str subregions.

These physiological domains provide a guide to *in vivo* stimulation location, to study regional functions of the Str, particularly in the CPu. As neighboring DA neurons engage in concerted burst activity (Beeler and Kisbye Dreyer, 2019; Parker et al., 2016), which extends to their terminals (Cai et al., 2020), activation of a subpopulation of terminals mimics physiologically relevant burst activity. So, the physiological domains mapped in this study may correspond to functional units controlled by concerted burst activity of DA neurons. Nonetheless, differences in the coherence of DA neuron activity is likely to differentially affect signal transduction. The Str domains identify representative recording locations necessary for the characterization of DA neuron transmission in the Str, in order to evaluate developmental, maturational and pathological changes. The domains may point to ways to achieve circuit specific pharmacotherapy by combinatorial drug targeting.

## Materials and Methods

### Mice

Mice were handled in accordance with the guidelines of the National Institutes of Health *Guide for the Care and Use of Laboratory Animals*, under protocol NYSPI-1494 approved by the Institutional Animal Care and Use Committee of New York State Psychiatric Institute. Mice were group housed and maintained on a 12 hour light/dark cycle. All slice/tissue preparations were done during the light phase. Food and water were supplied *ad libitum*. A total of 264 mice (124 male and 140 female), 2 to 3 months of age (postnatal days 60 to 91) were used.

Mice were C57BL6J/129Sv mixed background, backcrossed more than 5 times to C57BL6J and kept inbred. D2-EGFP mice, originally on a FVB background, were crossed to C57BL6J more than 8 times. DAT (Slc6a3)-internal ribosome entry site (IRES) cre (DAT^IREScre^) mice (Bäckman et al., 2006) (Jackson Laboratories, Bar Harbor, ME; RRID:IMSR_JAX:006660) were mated with ROSA26-floxSTOP-CAG-ChR2-EYFP (Ai32; ChR2-EYFP) (RRID:IMSR_JAX:024109) to generate mice with selective expression of ChR2 in DA neurons and enable selective stimulation of DA neuron terminals in the Str. Mice were homozygous for ChR2-EYFP to enable reliable stimulation. For identification of dSPN, iSPN, ChI, and FSI Str neurons, mice with fluorescent genetic markers for each neuron type, D1-tdTomato (RRID:IMSR_JAX:016204), D2-EGFP (GENSAT; RRID:MMRRC_000230-UNC), ChAT-EGFP (RRID:IMSR_JAX:007902), or PV-tdTomato (RRID:IMSR_JAX:027395), respectively, were bred with DAT^IREScre^;ChR2-EYFP double mutant mice. For recording from ChIs with K^+^-based pipette solution, double mutant of DAT^IREScre^; ChR2-EYFP mice without ChAT-EGFP were used, and ChIs were identified by soma size and membrane properties (Chuhma et al., 2014). D2-EGFP single mutant mice were used for immunohistochemical evaluation of D2-EGFP expression in ChIs.

### Brain slice electrophysiology

Mice were anesthetized with ketamine (90 mg/kg) /xylazine (7 mg/kg). After confirmation of full anesthesia, mice were decapitated and brains quickly removed in ice-cold high-glucose artificial cerebrospinal fluid (ACSF) (in mM: 75 NaCl, 2.5 KCl, 26 NaHCO_3_, 1.25 NaH_2_PO_4_, 0.7 CaCl_2_, 2 MgCl_2_ and 100 glucose, pH 7.4) saturated with carbogen (95% O_2_ and 5% CO_2_). Coronal 300 µm Str sections were cut with a vibrating microtome (VT1200S, Leica, Buffalo Grove, IL), allowed to recover in high glucose ACSF at room temperature for at least 1 hour, then transferred to the recording chamber (submerged, 500 µl volume) on the stage of an upright microscope (BX61WI, Olympus, Tokyo, Japan), continuously perfused with standard ACSF (in mM: 125 NaCl, 2.5 KCl, 25 NaHCO_3_, 1.25 NaH_2_PO_4_, 2 CaCl_2_, 1 MgCl_2_ and 25 glucose, pH 7.4) saturated with carbogen. ChR2-EYFP, D2-EGFP or ChAT-EGFP expression was confirmed by 470 nm LED field illumination; D1-tdTomato expression was confirmed by 530 nm LED illumination (DC4100, Thorlabs, Newton, NJ). Recorded neurons were visualized using enhanced visible light differential interference contrast (DIC) optics with a scientific c-MOS camera (ORCA-Flash4.0LT, Hamamatsu Photonics, Hamamatsu, Japan).

In DAT^IREScre^’;Ai32 mice, ChIs were identified visually by large soma size, confirmed by spontaneous firing, shallow resting membrane potentials (around −60 mV) and voltage sag with - 400 pA current injection (700 msec duration) (Chuhma et al., 2014). Recording patch pipettes (3-7 MΩ) were fabricated from standard-wall borosilicate glass capillary with filament (World Precision Instruments). Intracellular solution for DA PSC recording was (in mM): 135 K^+^-methane sulfonate (MeSO_4_), 5 KCl, 2 MgCl_2_, 0.1 CaCl_2_, 10 HEPES, 1 EGTA, 2 ATP and 0.1 GTP, pH 7.25. For glutamate and GABA PSC recording, Cs^+^-based pipette solution was used for better space clamp. For glutamate EPSC recording, K^+^-MeSO_4_ was replaced with Cs^+^-MeSO_4_, supplemented with QX314 (lidocaine N-ethyl bromide) 5 mM, to block action currents. For post-recording staining, Alexa Fluor 594 was added to the Cs^+^-MeSO_4_ based pipette solution. For GABA_A_ IPSC recording, K^+^-MeSO_4_ was replaced with CsCl supplemented with QX314. All recordings were done under whole cell voltage clamp at −70 mV with an Axopatch 200B amplifier (Molecular Devices, San Jose, CA). Series resistance (8-30 MΩ) was compensated online by 70-80%. Liquid junction potentials (5-12 mV) were adjusted online. Recording began 2 min after entering whole cell mode, to allow for diffusion of pipette solution. Synaptic responses were evoked with 5 msec field illumination with a high-power blue LED (470 nm; Thorlabs) delivered as a single pulse at 0.1 Hz with maximum intensity (10 V) and full field illumination, through a 60x (NA 0.90) water-immersion objective (Olympus) for consistent activation of DA neuron terminals in the field of view.

DA neuron PSCs were pharmacologically isolated by blocking all other transmitter receptors, e.g. co-transmitted glutamate PSCs were isolated by blocking D1/5, D2, and GABA receptors. The antagonist-isolation cocktail was applied at least 12 min prior to recording. Nicotinic and muscarinic ACh receptors were continuously blocked as well, since even in slices ChIs are spontaneously active and there is a basal cholinergic tone, that is likely to show regional heterogeneity. Antagonists and their concentrations were: iGluR AMPA/kainate CNQX 20 µM, iGluR NMDA D-APV 50 µM, mGluR1 JNJ16259685 5 µM, D1/5R SCH23390 10 µM, D2 S(-)- sulpiride 10 µM, GABA_A_R SR95531 (gabazine) 10 µM, GABA_B_R CGP55845 3 µM, nAChR mechamilamine 10 µM, mAChR scopolamine 2 µM. To achieve faster blockade of PSCs for confirmation of isolated responses (Extended Data Fig. 2), doubled concentration CNQX, D-APV, or JNJ was used. Stock solutions of drugs were prepared in either water or DMSO, and diluted by 1000 to 5000 fold in recording ACSF. Drugs were applied by perfusion. Locations of recorded cells were mapped on mouse atlas coronal sections (Paxinos and Franklin, 2008), on a 50 µm medial-lateral and dorsal-ventral grid.

Voltage clamp data were filtered at 5 kHz with a 4-pole Bessel filter, digitized at 5 kHz (Digidata 1550A, Molecular Devices), recorded using pClamp 10 (Molecular Devices; RRID:SCR_011323), and analysed with Axograph X (Axograph Science; RRID:SCR_014284). PSCs were measured from averages of 10 consecutive traces. Synaptic responses were measured starting with the second stimulus to avoid artificially large responses to the first stimulus elicited after a period of rest. Because of synchronized and fast responses, iGluR and GABA_A_ PSCs were evaluated by peak amplitude measured in a 15 msec post-stimulus window. mGluR, D2R, and D1/5R PSCs in ChIs were evaluated by charge transfer measured in a 1.5 sec post-stimulus window, as area under the curve. D1/5R PSCs in FSIs were evaluated in a 0.3 sec window due to their shorter time course (slower than PSCs in ChIs), and because they were not as synchronized (as glutamate or GABA_A_R PSCs). GABA_A_ PSCs declined quickly with the first 10 stimuli (Straub et al., 2014), so we measured GABA_A_ PSC amplitudes in averages of responses to the second 10 stimuli. Rise time (10-90% peak) and width at 50% peak of D1/5R PSCs were measured in ChIs with responses greater than 2 pC and FSIs with peak amplitudes greater than 8 pA. Baseline charge transfer was measured from 10 consecutive traces without photostimulation. For comparison between ChIs identified by ChAT-EGFP fluorescence and ChIs identified by post-recording staining, we chose three discrete locations and compared PSC measures. The three locations were NAc medial shell (in NAc shell region in mouse atlas, and ML ≤ +1; in mm from bregma), medial caudate putamen (CPu) (AP ≥ +0.26 and ≤ +1.1, ML ≥ +2.1, DV ≥ –3.8), and lateral CPu (AP ≥ +0.95 and ≤ +1.34, ML ≤ +1.15, DV ≥ –3.7). In total 1677 cells were used for this study, and among them, 34 were the same cells reported in Chuhma et al. (2018). A maximum of 10 cells were recorded per animal.

### Immunocytochemistry

Three D2-EGFP mice (P74, 1 male and 2 females) were anesthetized with ketamine/xylazine and perfused with cold phosphate buffered saline (PBS) followed by 4% paraformaldehyde (PFA). Brains were removed and post-fixed overnight in 4% PFA. Coronal 50 µm sections were cut using a vibrating microtome (Leica VT1200S), and stored in a cryoprotectant solution (30% glycerol, 30 % ethylene glycol in 0.1 M Tris HCl pH 7.4) at −20 °C until processing. Sections were washed in PBS and incubated in glycine (100 mM) for 30 min to quench aldehydes. Non-specific binding was blocked with 10% normal goat serum (NGS; Millipore, St. Louis, MO) in 0.1 PBS Triton X-100 (PBS-T) for 2 hours. Primary antibodies in 0.02% PBS-T and 2 % NGS were applied for 24 hours, at 4 °C on a shaker. Primary antibodies were: anti-EGFP (1:2,000 dilution, rabbit polyclonal, Millipore, RRID:AB_91337) and anti-choline acetyltransferase (ChAT; 1:1000 dilution, goat polyclonal, Millipore, RRID:AB_2079751). For post-recording staining, slices were fixed with 4% PFA at room temperature for 1-3 hours. After washing with PBS, slices were processed as above, except that the primary antibody (anti ChAT) was applied for 2-3 days. The recorded cells were observed directly with Alexa Fluor 594 dye fluorescence without immunostaining. Sections were then washed with PBS and secondary antibodies applied for 45 min in 0.02% PBS-T at room temperature. Secondary antibodies (1:200 dilution; ThermoFisher Scientific, Waltham, MA) were: anti-goat Alexa Fluor 488 (RRID:AB_2534102), anti-goat Alexa Fluor 594 (RRID:AB_2534105) and anti-rabbit Alexa Fluor 488 (RRID:AB_2535792). Sections were mounted on gelatin subbed slides (Southern Biotech, Birmingham, AL) and cover slipped with Prolong Gold aqueous medium (ThermoFisher Scientific) and stored at 4 °C.

### Imaging and cell counting

Images were acquired with an AxioImager.M2 fluorescence microscope and ZEN Digital Imaging software (Zeiss; RRID:SCR_013672) using a 20x objective for GFP/ChAT colocalization and a 40x objective for Alexa 594/ChAT colocalization and dendrite tracing. Images were taken in 1 µm steps subtending the entire slice thickness (26-37 images per slice), and each z-section image was examined for immuno- or dye-fluorescence. ChAT^+^ and ChAT^+^/GFP^+^ cells were counted using the Optical Fractionator Probe in Stereo Investigator 11 (MBF Bioscience, Williston, VT; RRID:SCR_002526) using a 10x objective, over 20% of the area of every 5th slice. In each brain, 12-15 slices were analyzed. Stereological studies were performed unilaterally.

EGFP fluorescence intensity and soma size measurements in D2-EGFP mouse slices were measured with Image J (FIJI) 2.0.0 (NIH, RRID:SCR_003070). Regions of interest (ROIs) were made by manual tracing of ChAT^+^ or GFP^+^ somas, and the area used as a measure of soma size. Average GFP staining intensity in each ROI was regarded as GFP intensity of each cell. For ChAT^+^/GFP^+^ cells, ROIs made in ChAT channel projected to the GFP channel and measured averaged GFP staining intensity within those.

Dendrites of Alexa-filled ChIs were traced with Image J 2.0.0 using Simple Neurite Tracer plug-in. The same set of cells used for post-recording ChAT staining were used for dendrite tracing, excluding one cell with faint dendrite staining. Tracing was done in Alexa 594 z-stack images. For visualization of fine dendrites, contrast was enhanced. Dendrites were semi-automatically traced until dendrites became not recognizable or reached a cut end. Generally, tracing extended to the tertiary dendrites. Lengths of all measured dendrites of individual cells were summed.

### Statistics for electrophysiology and immunostaining

Statistics, including sample size estimation, were done in R3.6.1 (RRID:SCR_001905) with RStudio 1.2.1335 (RRID:SCR_000432). The following packages were used in addition to base/stats; pwr 1.2-2 (for sample size estimation), WebPower 0.5.2 (for sample size estimation), afex 0.25-1 (for Type III sum of square ANOVA), emmeans 1.4.1 (for post-hoc comparison). We used parametric tests, as the variables we measured were continuous numeric variables (not ranked variables).

Sample sizes were estimated with power = 0.8 and alpha = 0.05. Effect sizes were calculated and estimated from previously performed similar experiments. For paired t-test (antagonist effects), Cohen’s d was set to 1.6 (very strong effect), suggesting 6 cells (pairs) were required. For one-way ANOVA for image analysis, Cohen’s f was set to 0.5 (strong effect), suggesting that a total 42 cells, in 3 groups. For two-way ANOVA, Cohen’s f was set to 0.6 based on a previously performed regional heterogeneity study, giving a sample size of 30 cells, in 6 groups. For the correlation study, a significance criterion of R^2^ = 0.5 (R = 0.71) gave 13 as the required number of observations.

Data are presented as mean ± S.E.M, unless otherwise noted. Kormogorov-Smirnov tests were one-tailed, while all other tests were two-tailed. The test used, t value, F value, R^2^, degree of freedom, p value, or exact number of samples (n), and what n stands for are reported in either the main text or figure legends. For antagonist effects on pharmacologically isolated PSCs (Fig. 2), we include a separate table with statistical results (Table S1). Significance was set at p < 0.05.

### Spatial data analysis

All spatial data analyses were done in R3.6.1. The following packages were used for data analysis and plotting, in addition base/stats; dplyr 0.8.1 (RRID:SCR_016708), tidyr 0.8.3 (RRID:SCR_003005), purrr 0.3.2, readr 1.3.1, readxl 1.3.1, ggplot2 3.1.1 (RRID:SCR_014601), RColorBrewer 1.1-2 (RRID:SCR_016697), gridExtra 2.3, egg 0.4.2, cowplot 0.9.4, plotly 4.9.0 (RRID:SCR_013991), sp 1.3-1, gstat 2.0-2, factoextra 1.0.5 (RRID:SCR_016692), and ClustGeo 2.0.

PSC sizes were estimated on a 150 µm grid, manually overlaid on coronal sections (∼150 µm apart) of a mouse brain atlas (Paxinos and Franklin, 2008). In the unilateral Str, the number of grid lattices was 4177. Estimation was done by ordinary block kriging with a 450 µm cube block with gstat 2.0-2 (Gräler et al., 2016). Experimental variograms were fit with Matern covariance functions, and parameters (psill, nugget, and range) of each map are reported in Table S2.

Hierarchical clustering of kriging results was done with spatial constraint with ClustGeo 2.0 (Chavent et al., 2018). Estimated PSC sizes were normalized by the maximum and minimum values of each map before clustering. Optimal numbers of clusters (K) were chosen by the silhouette method with factoextra 1.0.5 in 3-10 clusters to obtain contiguous subdomains. After an optimal K was obtained, mean and SD of PSCs in each cluster were calculated, and compared to the results among K and K ± 1 clusters, to confirm that the K was optimal. If there were more than two clusters with almost the same set of mean values, K was likely to be too large (too many clusters). If adding one more cluster largely reduced SD in clusters, K was likely too small. The chosen K was optimal in cell-type clusters, while K+1 was optimal for transmitter clusters. Location and data (PSC size) dissimilarity matrices were calculated using the Euclidean distances of values between data points.

To obtain the optimal mixing parameter alpha, a quality criterion (Q) was calculated in 0.1 steps with the ClustGeo 2.0 *choicealpha* function (Chavent et al., 2018). Alpha ranges from 0, where clustering is determined solely by location, to 1, where clustering is determined solely by data (PSC size). With increasing alpha, the data matrix contribution is reduced, while the location matrix contribution is increased. Generally, reduction of the data matrix-contribution caused reduced homogeneity of data in the clusters (reducing data Q), while geographic (location) coherence increased (increasing location Q). Alpha is optimized when location Q increases significantly, while minimizing data Q reduction. However, our datasets were distributed in three dimensions and we needed stronger location constraint than the ward-wise socio-demographic analysis for which the ClustGeo package was developed (Chavent et al., 2018), to obtain contiguous subdomains. Therefore, we used the alpha value just after location and data normalized Q plot curves crossed. The determined alphas were: dSPN 0.3, iSPN 0.3, ChI 0.4, FSI 0.4, glutamate 0.3, DA 0.4, and GABA 0.3. After clustering with the determined alpha, dendrograms were cut at the points to obtain optimal number of clusters. Means and SDs of transmitter receptor/cell type responses in each cluster were calculated. We used the number of grid points in each cluster to estimate the volume of each cluster. To evaluate the quality of clustering, F values were calculated for each variable. F is defined as between-cluster variance divided by within-cluster variance, and magnitudes of F values indicate how well the respective variables discriminate between clusters (Calinski and Harabasz, 1974).

## Acknowledgements

We thank Casey Brody and Sarah Garcia for technical assistance. This work was supported by NIH R01 DA038966 and R01 MH117128 (SR).

---

**Supplementary Fig 1.**
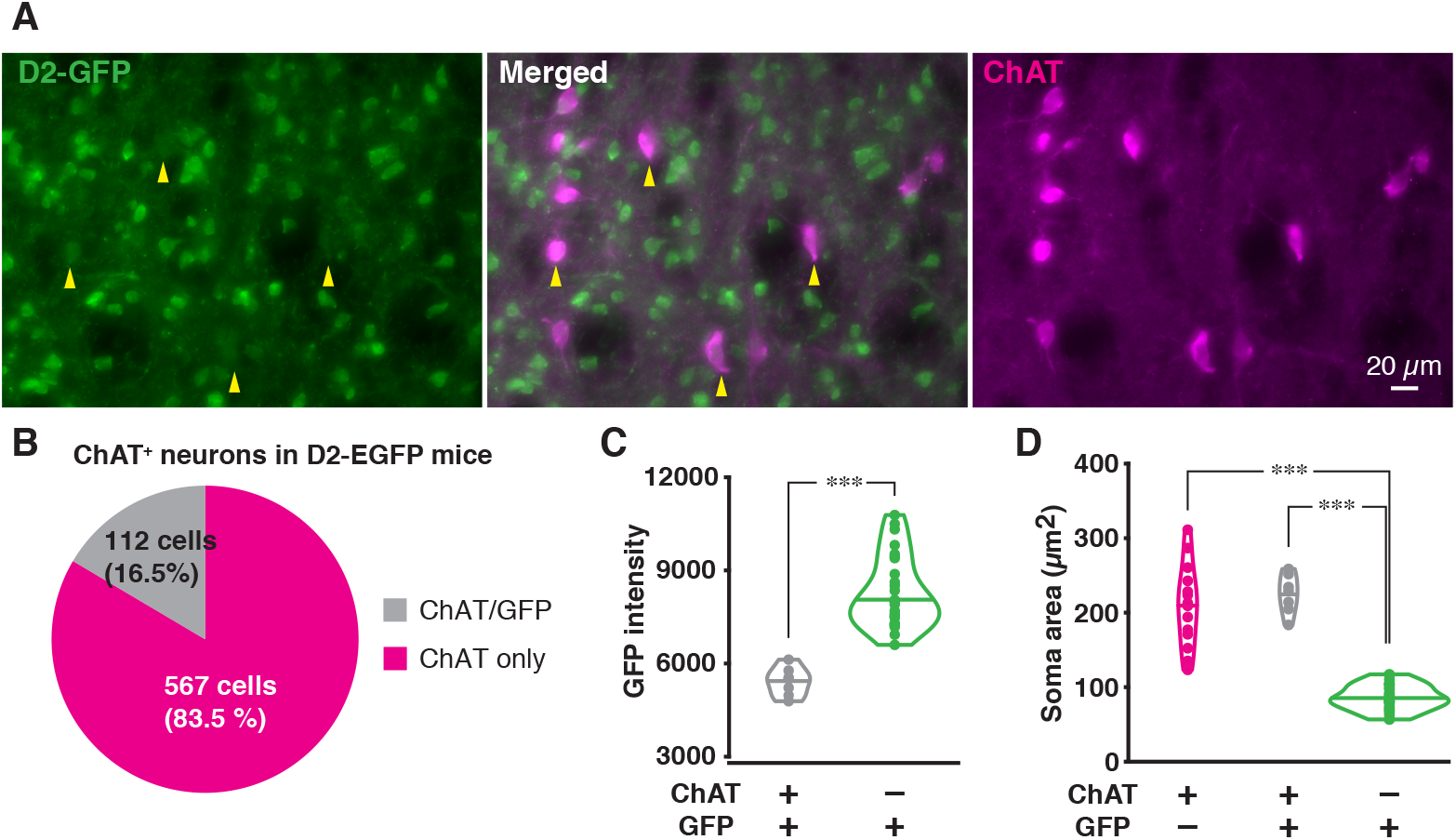
EGFP expression in D2-EGFP mice. In slices from D2-EGFP mice, brighter EGFP^+^ neurons were iSPNs and not ChIs. (**A**) Representative EGFP (green) and ChAT (magenta) immunostaining in the Str of a D2-EGFP mouse. Yellow arrows indicate EGFP^+^/ChAT^+^ cells. (**B**) The ratio of EGFP^+^ (gray) and EGFP^−^ (magenta) ChAT^+^ neurons. Cell numbers are stereological counts from three mice. (**C**) EGFP fluorescence intensity in ChAT^+^ (gray) and ChAT^−^ (green) neurons. EGFP fluorescence in EGFP^+^/ChAT^+^ neurons was dimmer than in EGFP^+^/ChAT^−^ neurons. Each dot shows the average EGFP fluorescence intensity of a single soma. Outlines in the graph show densities of counts and horizontal lines in the outlines show medians of the intensities. EGFP^+^/ChAT^+^ n = 7 cells, EGFP^+^/ChAT^−^ n = 28 cells, Welch’s t-test, t = −10.3, df = 25.2, p < 0.0001. (**D**) Soma size analysis of EGFP^−^/ChAT^+^ (magenta), EGFP^+^/ChAT^+^ (gray), and EGFP^+^/ChAT^−^ (green) neurons. ChAT^+^ neurons, regardless of EGFP expression, were significantly larger than ChAT^−^ neurons. EGFP^−^/ChAT^+^ n = 14 cells, one-way ANOVA, F(2, 46) = 88.8, p < 0.0001. Pairwise comparison as post-hoc test (with Bonferroni correction) EGFP^−^/ChAT^+^ – EGFP^+^/ChAT^+^, p = 0.92; EGFP^−^/ChAT^+^ – EGFP^+^/ChAT^−^, p < 0.0001; EGFP^+^/ChAT^+^ – EGFP^+^/ChAT^−^, p < 0.0001.

**Supplemental Fig 2.**
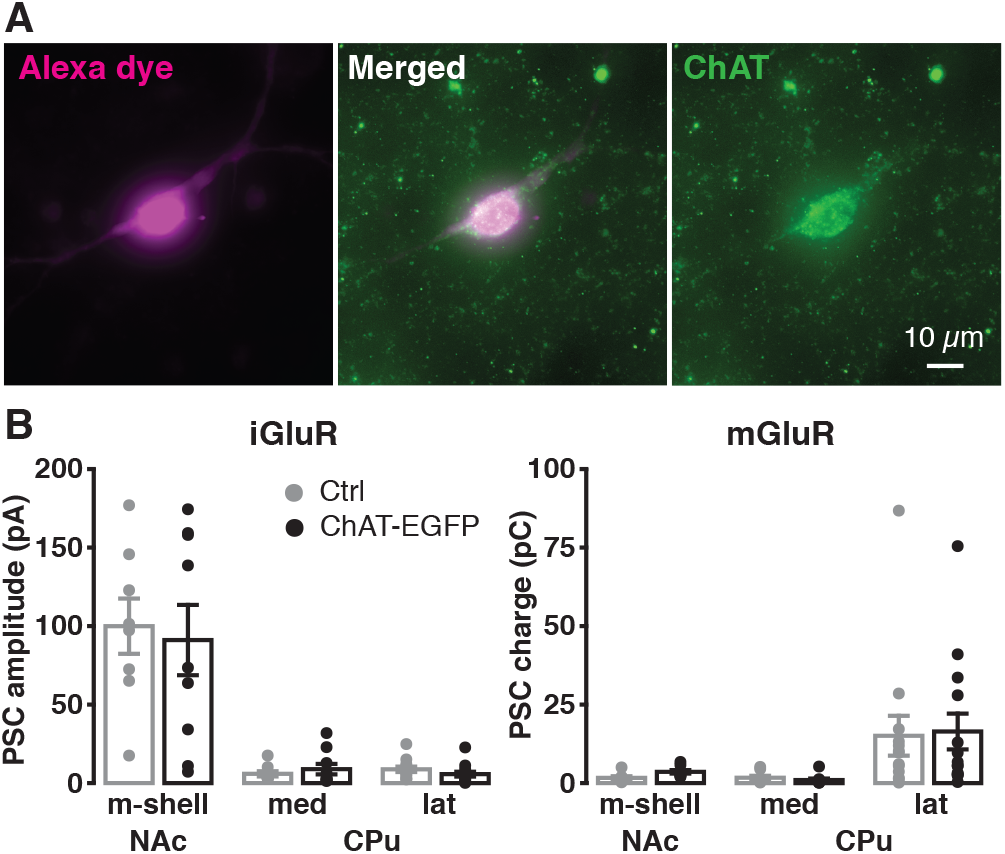
Glutamate responses in ChAT-EGFP mice and mice without ChAT-EGFP. VAChT overexpression in ChAT-EGFP mice did not affect measures of regional heterogeneity. The identification of ChIs in mice without ChAT-EGFP (DAT^IREScre/+^; ChR2) was confirmed by post-recording ChAT immunostaining. (**A**) Images of a recorded ChI filled with Alexa Fluor 594 (magenta), ChAT counter staining of the cells (green), and a merged image (white). Note that the green channel also includes DA neuron terminals with ChR2-EYFP fluorescence. (**B**) iGluR (left) and mGluR (right) PSCs sizes on ChIs in NAc medial shell (m-shell), medial caudate-putamen (med CPu), and lateral (lat) CPu from mice without ChAT-EGFP (Ctrl; gray) and with ChAT-EGFP (black). There were no genotypic differences. Recorded number of cells were: NAc m-shell Ctrl n = 8 cells, ChAT-EGFP n = 9 cells; med CPu Ctrl n = 9 cells, ChAT-EGFP n = 10 cells; lat CPu Ctrl n = 13 cells, ChAT-EGFP n =14 cells. iGluR: location F(2, 57) = 49.5, p < 0.0001; genotype F(1, 57) = 0.14, p = 0.71; interaction F(2, 57) = 0.16, p = 0.85; two-way ANOVA. mGluR: location F(2, 57) = 6.8, p = 0.002, genotype F(1, 57) = 0.05, p = 0.82, interaction F(2, 57) = 0.04, p = 0.96; two-way ANOVA.

**Supplemental Fig 3.**
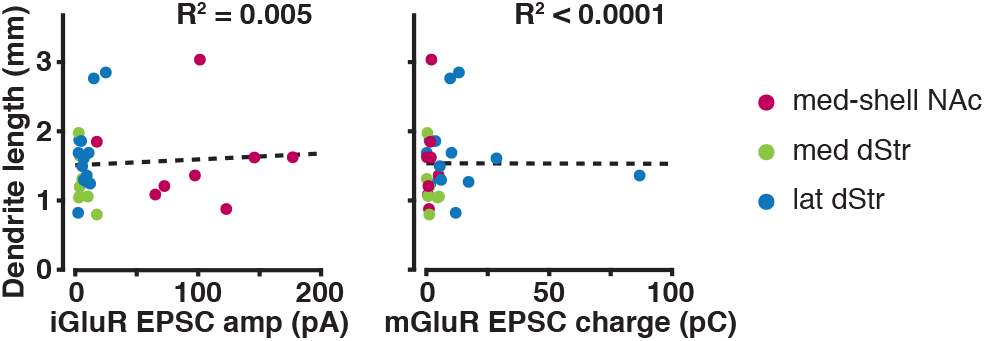
Correlation between dendrite lengths and PSC sizes. Regional differences in synaptic responses were not due to regional differences in the truncation of dendrites by slicing. The same set of cells used in Supplemental Fig 2 were analyzed. One cell did not show good dendrite staining and was discarded. Dendrite tracing was done in Alexa Fluor images with contrast enhancement. Total lengths of traceable dendrites (in mm) were plotted against amplitude of iGluR PSCs (left) or charge transfer of mGluR PSCs (right). Locations of recorded cells are color coded: NAc m-shell, dark red; med CPu, light green; lat CPu, blue. Smaller responses did not correlate with shorter dendrites. iGluR: R^2^ = 0.005, F(1,27) = 0.145, p = 0.71. mGluR: R^2^ < 0.0001, F(1,27) = 0.0001, p = 0.99.

**Supplemental Fig 4.**
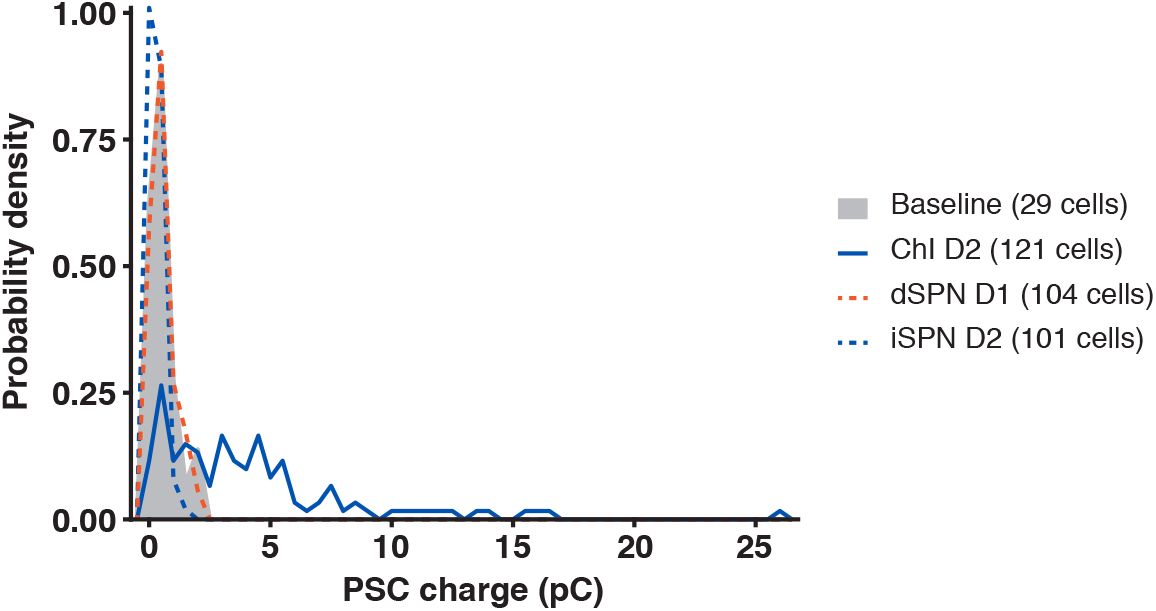
No significant DA responses in dSPNs or iSPNs. D1/5R PSC charge transfer in dSPNs (orange broken line) and D2R PSC charge transfer in iSPNs (blue broken line) were compared to baseline (gray shading). For baseline, charge transfers were measured in a 1.5 sec time window without photostimulation. The charge transfer histograms (bin width 0.5 pC) were expressed as a probability density, and outlines of the histograms are shown. For comparison, the probability density curve of the positive response of D2R PSCs in ChI is shown (blue solid line). Comparison to baseline (n = 29 cells): D1/5R PSC in dSPN, n = 104 cells, D = 0.13, p = 0.48; D2R PSC in iSPN, n = 101 cells, D = 0.14, p = 0.42; D2R PSC in ChI, n = 121 cells, D = 0.65, p < 0.0001; one-tailed two-sample Kolmogorov-Smirnov test.

**Supplemental Fig 5.**
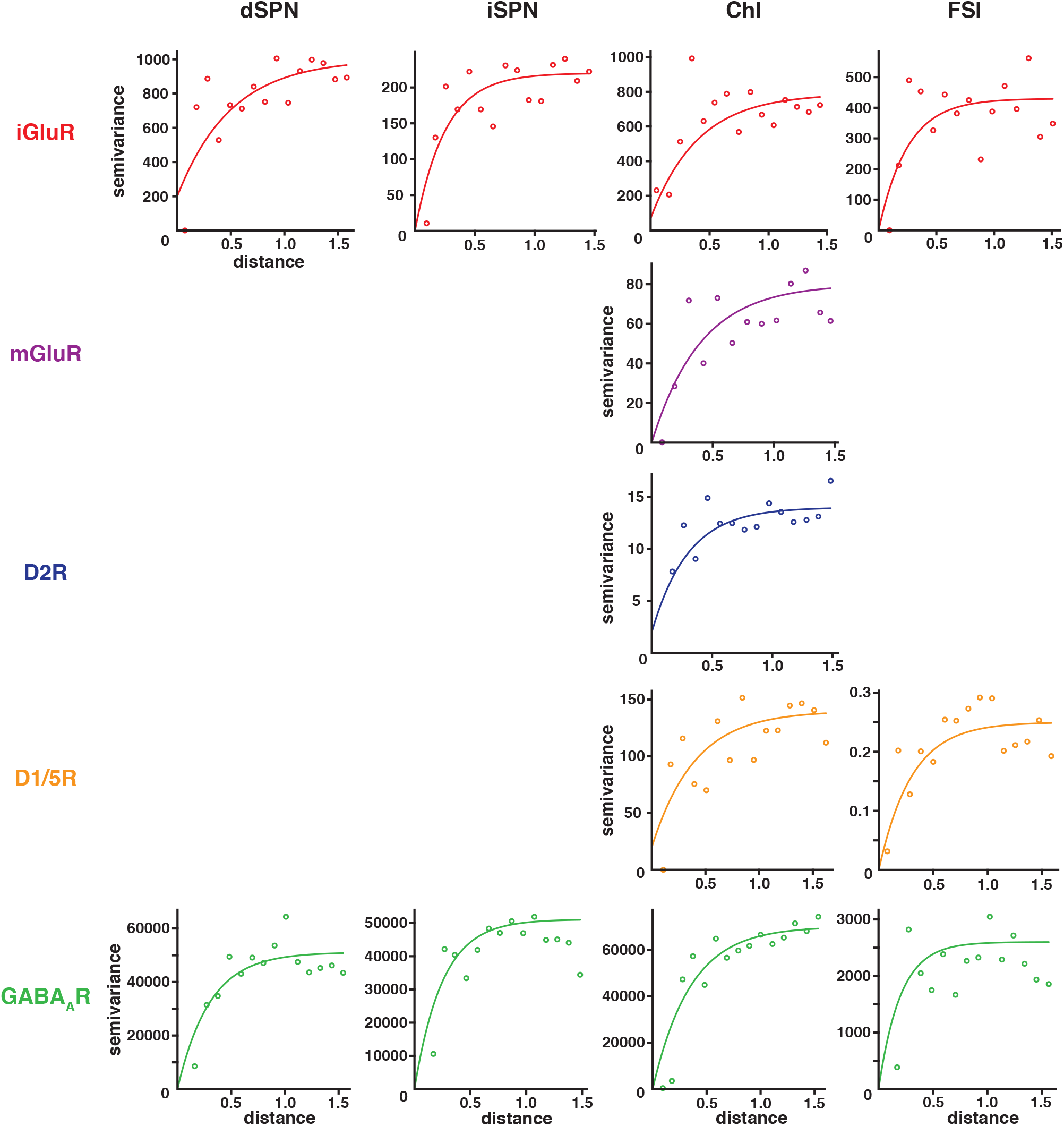
Experimental variograms of recorded PSCs. Semivariance of PSCs in pairs of recording locations (vertical axis) were plotted versus euclidean distance of these locations (in mm). Averages in 0.1 mm distance bins are shown as open circles in the graphs. Variograms were fit by Matern covariance functions (lines), and the fitted values were used to obtain weights for averaging to estimate PSC sizes on a cubic grid. See Supplemental Table 2 for fitting parameters.

**Supplemental Table 1.** Paired t-test for antagonist effects shown in Fig 2.

|  | iGluR |  |  |  | mGluR | D2R | D1/5R |  | GABA<sub>A</sub>R |  |  |  |
| --- | --- | --- | --- | --- | --- | --- | --- | --- | --- | --- | --- | --- |
|  | dSPN | iSPN | ChI | FSI | ChI | ChI | ChI | FSI | dSPN | iSPN | ChI | FSI |
| **df** | 8 | 7 | 7 | 7 | 3 | 6 | 6 | 7 | 8 | 8 | 8 | 7 |
| **t** | 6.9 | 3.4 | 3.1 | 3.8 | 3.7 | 3.7 | 7.4 | 4.5 | 3.0 | 2.5 | 3.8 | 8.5 |
| **p** | 0.0001 | 0.011 | 0.017 | 0.0063 | 0.034 | 0.0099 | 0.0003 | 0.0029 | 0.017 | 0.035 | 0.005 | 0.01 |

**Supplemental Table 2.**
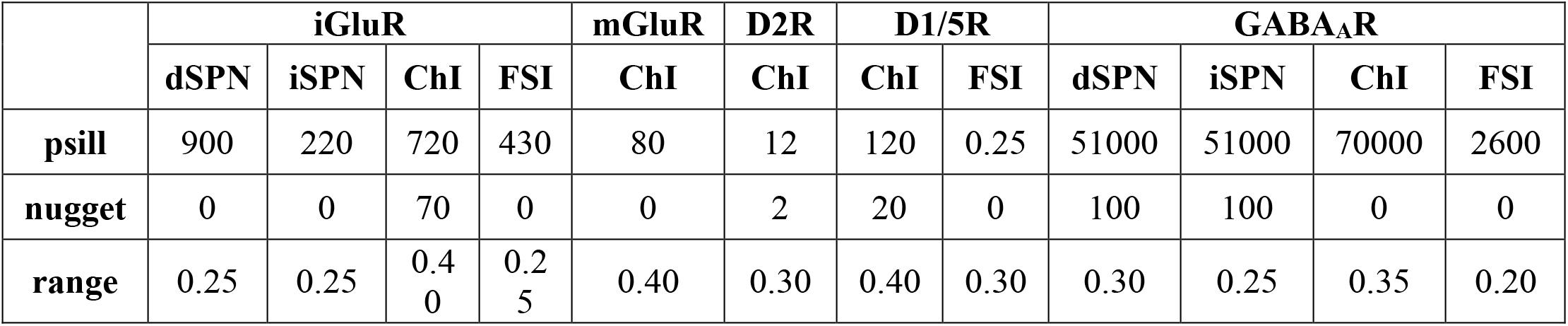
Experimental variogram fitting parameters.

|  | iGluR |  |  |  | mGluR | D2R | D1/5R |  | GABA<sub>A</sub>R |  |  |  |
| --- | --- | --- | --- | --- | --- | --- | --- | --- | --- | --- | --- | --- |
|  | dSPN | iSPN | ChI | FSI | ChI | ChI | ChI | FSI | dSPN | iSPN | ChI | FSI |
| psill | 900 | 220 | 720 | 430 | 80 | 12 | 120 | 0.25 | 51000 | 51000 | 70000 | 2600 |
| nugget | 0 | 0 | 70 | 0 | 0 | 2 | 20 | 0 | 100 | 100 | 0 | 0 |
| range | 0.25 | 0.25 | 0.4<br>0 | 0.2<br>5 | 0.40 | 0.30 | 0.40 | 0.30 | 0.30 | 0.25 | 0.35 | 0.20 |

**Supplemental Table 3.** Descriptive statistics of cell-type clusters and anatomical locations

| dSPN | Clust. 1 | Clust. 2 | Clust. 3 | F |
| --- | --- | --- | --- | --- |
| Location<sup>1</sup> | Med. NAc<br>Ant.Lat.Dors. CPu | Lat. NAc<br>Ant. CPu | Post. CPu |  |
| N of grid points | 721 | 2011 | 1445 |  |
| iGluR (pA) mean<br>*SD* | 42.3<br>13.6 | 13.7<br>5.8 | 16.1<br>4.3 | 4137 |
| GABA<sub>A</sub> (pA) mean<br>*SD* | 208.3<br>83.9 | 194.6<br>93.8 | 106.9<br>45.4 | 647 |

| iSPN | Clust. 1 | Clust. 2 | Clust. 3 | F |
| --- | --- | --- | --- | --- |
| Location | Med. NAc<br>Ant.Lat.Dors. CPu | Ant.Med. CPu | Lat. NAc<br>Post. CPu |  |
| N of grid points | 672 | 958 | 2547 |  |
| iGluR (pA) mean<br>*SD* | 17.9<br>6.7 | 7.2<br>3.7 | 8.4<br>3.7 | 1459 |
| GABA<sub>A</sub> (pA) mean<br>*SD* | 135.8<br>59.5 | 213.6<br>76.9 | 87.9<br>37.3 | 2001 |

| ChI | Clust. 1 | Clust. 2 | Clust. 3 | Clust. 4 | Clust. 5 | Clust. 6 | F |
| --- | --- | --- | --- | --- | --- | --- | --- |
| Location | NAc | Ant. CPu | Med. CPu | Lat. CPu | Dors. CPu | Post. CPu |  |
| N of grid points | 394 | 1227 | 447 | 679 | 685 | 745 |  |
| iGluR (pA) mean<br>*SD* | **65.8**<sup>2</sup><br>25.1 | 17.5<br>12.3 | 19.2<br>8.9 | 11.7<br>5.2 | 12.9<br>4.3 | 12.4<br>4.3 | 1533 |
| mGluR (pC) mean<br>*SD* | 2.9<br>0.8 | 3.1<br>1.6 | 2.5<br>0.7 | **10.3**<br>6.8 | 2.6<br>1.3 | 1.8<br>0.7 | 773 |
| D2R (pC) mean<br>*SD* | 3.0<br>0.8 | 4.1<br>0.9 | **7.4**<br>2.2 | 3.4<br>0.7 | 4.8<br>1.0 | 3.5<br>0.5 | 1119 |
| D1/5R (pC) mean<br>*SD* | 2.8<br>0.6 | 5.9<br>3.6 | 6.8<br>3.4 | **12.3**<br>5.3 | 10.0<br>5.0 | 4.4<br>2.1 | 578 |
| GABA<sub>A</sub> (pA) mean<br>*SD* | 142.1<br>47.3 | 284.1<br>79.7 | **384.2**<br>93.5 | 187.7<br>77.8 | 163.5<br>55.8 | 122.4<br>72.9 | 1079 |

| FSI | Clust. 1 | Clust. 2 | Clust. 3 | Clust. 4 | Clust. 5 | F |
| --- | --- | --- | --- | --- | --- | --- |
| Location | Vent. NAc | Ant. CPu | Ant.Med. CPu | Med.Dors. CPu<br>Ant.Dors.Lat CPu | Post. CPu |  |
| N of grid points | 646 | 1628 | 303 | 229 | 1371 |  |
| iGluR (pA) mean<br>*SD* | 10.1<br>3.7 | 8.4<br>2.9 | 6.6<br>2.3 | 29.2<br>10.4 | 12.1<br>4.5 | 1285 |
| D1/5R (pC) mean<br>*SD* | 0.26<br>0.10 | 0.50<br>0.16 | 0.98<br>0.17 | 0.41<br>0.10 | 0.28<br>0.09 | 2180 |
| GABA<sub>A</sub> (pA) mean<br>*SD* | 24.6<br>5.1 | 35.6<br>12.8 | 32.1<br>6.9 | 46.1<br>10.6 | 23.5<br>5.9 | 524 |

**Supplemental Table 4.** Descriptive statistics of transmitter clusters and anatomical locations

| Glutamate | Clust. 1 | Clust. 2 | Clust. 3 | Clust. 4 | F |
| --- | --- | --- | --- | --- | --- |
| Location | NAc | Ant.Med. CPu<br>Ant.Lat.Vent. CPu | Ant.Lat.Dors. CPu | Post. CPu |  |
| N of grid points | 417 | 1777 | 746 | 1237 |  |
| dSPN iGluR (pA) mean<br>*SD* | 40.1<br>14.8 | 12.5<br>4.5 | 30.9<br>14.8 | 15.7<br>4.1 | 1697 |
| iSPN iGluR (pA) mean<br>*SD* | 17.2<br>5.4 | 7.3<br>4.7 | 13.6<br>6.4 | 8.1<br>1.9 | 780 |
| ChI iGluR (pA) mean<br>*SD* | 66.7<br>23.5 | 17.2<br>9.3 | 9.1<br>3.5 | 13.7<br>4.7 | 3504 |
| FSI iGluR (pA) mean<br>*SD* | 15.5<br>9.4 | 7.3<br>2.5 | 15.4<br>9.1 | 11.7<br>3.3 | 578 |
| ChI mGluR (pC) mean<br>*SD* | 3.0<br>0.7 | 2.9<br>1.6 | 9.5<br>6.8 | 2.1<br>1.0 | 1006 |

| Dopamine | Clust. 1 | Clust. 2 | Clust. 3 | Clust. 4 | F |
| --- | --- | --- | --- | --- | --- |
| Location | NAc | Ant.Med. CPu | Ant.Lat.Dors. CPu | Post. CPu |  |
| N of grid points | 904 | 938 | 1210 | 1125 |  |
| ChI D2R (pC) mean<br>*SD* | 3.4<br>0.8 | 5.9<br>2.2 | 4.0<br>1.1 | 3.8<br>0.8 | 657 |
| ChI D1/5R (pC) mean<br>*SD* | 3.7<br>2.3 | 5.2<br>2.2 | 12.6<br>5.1 | 5.4<br>2.7 | 1533 |
| FSI D1/5R (pC) mean<br>*SD* | 0.28<br>0.09 | 0.72<br>0.23 | 0.38<br>0.17 | 0.34<br>0.11 | 1525 |

| GABA | Clust. 1 | Clust. 2 | Clust. 3 | Clust. 4 | F |
| --- | --- | --- | --- | --- | --- |
| Location | NAc<br>Ant.Vent. CPu | Ant.Med.Dors. CPu | Ant.Lat.Dors. CPu | Post. CPu |  |
| N of grid points | 1177 | 701 | 858 | 1441 |  |
| dSPN (pA) mean<br>*SD* | 145.2<br>58.8 | 265.7<br>98.9 | 217.4<br>75.1 | 105.8<br>45.4 | 1111 |
| iSPN (pA) mean<br>*SD* | 106.2<br>54.8 | 188.7<br>68.5 | 177.1<br>79.1 | 76.8<br>32.7 | 907 |
| ChI (pA) mean<br>*SD* | 235.9<br>103.0 | 357.4<br>89.0 | 158.6<br>63.8 | 168.4<br>83.8 | 904 |
| FSI (pC) mean<br>*SD* | 32.1<br>14.5 | 30.9<br>8.2 | 37.1<br>11.4 | 24.4<br>6.3 | 280 |

